# ECSIT prevents Alzheimer’s disease pathology by regulating neuronal mitochondrial ROS and mitophagy

**DOI:** 10.1101/2020.01.09.900407

**Authors:** Alice Lepelley, Lauren S. Vaughn, Agnieszka Staniszewski, Hong Zhang, Fang Du, Peter Koppensteiner, Zeljko Tomljanovic, Flávia R.G. Carneiro, Andrew F. Teich, Ipe Ninan, Shirley ShiDu Yan, Thomas S. Postler, Matthew S. Hayden, Ottavio Arancio, Sankar Ghosh

## Abstract

Altered mitochondrial fitness is a potential triggering factor in Alzheimer’s disease (AD). Mitochondrial quality control pathways are dysfunctional and mitochondrially-derived reactive oxygen species (mROS) levels are increased in AD patient brains. However, the pathways responsible for dysregulated mROS accumulation have remained relatively unclear. In this study, we demonstrate that levels of ECSIT, a mitochondrial oxidative phosphorylation (OxPhos) complex I (CI)-associated protein, are reduced in AD-affected brains. Neuronal ECSIT downregulation increased mROS generation and impaired mitophagy of defective mitochondria. Consequently, decreasing neuronal ECSIT caused AD-like changes, including memory loss and neuropathology. In contrast, augmented neuronal expression of ECSIT protected against the development of an AD-like phenotype. Decreased levels of ECSIT in AD patient brains therefore likely contribute to oxidative stress, neuroinflammation and AD pathogenesis.

## Main

Reactive oxygen species (ROS) are believed to be a major cause of cellular damage during aging and contribute to age-related progressive neuropathologies, including Alzheimer’s disease (AD)^1^. Early, widespread oxidative stress is observed in AD and is postulated to be a key driver of neurodegeneration^2^. However, the source of this oxidative stress has been elusive. Neuronal metabolism relies heavily on mitochondrial oxidative phosphorylation (OxPhos)^3^. The mitochondrial respiratory chain is a major source of ROS (mROS) in the central nervous system (CNS) under normal physiological conditions. When mitochondrial function is compromised, oxidative stress is increased^4^. Mitochondrial dysfunction, as evidenced by downregulation of mitochondrial respiratory chain complexes and altered respiration, is one of the earliest and most prominent features in the brain of AD patients, and has been hypothesized to be the source of oxidative stress in AD^5–8^. Although mROS are unstable and difficult to measure in AD patients, studies have shown evidence of mROS accumulation through oxidative modifications of proteins and lipids^9^ and increased oxidative stress in cells reconstituted with mitochondria isolated from AD patients^10^. Moreover, ROS accumulation and mitochondrial dysfunction in animal models of AD have implicated mROS in disease pathogenesis^11–15^, while scavenging mROS can ameliorate neuropathology^4^. However, despite the evidence supporting a pathogenic role for mROS, mechanistic insights into the etiology of mROS accumulation in AD are still lacking^16^.

Normally, mROS production is limited by removal of dysfunctional mitochondria through mitophagy, a selective autophagy pathway targeting mitochondria^17^. Autophagy is an evolutionarily conserved process during which cellular components are engulfed in autophagic vesicles, fused to lysosomes and degraded. Autophagy is necessary for cellular viability, and autophagic dysfunction has been linked to neurodegeneration, aging, and age-related diseases^18^. Mitophagy specifically has been proposed to contribute to several neurodegenerative diseases^18^. Evidence of early mitophagy induction and dysfunction has been observed in AD brains and in cells derived from patients with familial and sporadic forms of AD, which may exacerbate mitochondrial oxidative stress^12, 19–22^.

ECSIT was originally identified and characterized as a signaling adapter in innate immune pathways^23, 24^. Subsequent work demonstrated that ECSIT localizes to mitochondria, where it interacts with the assembly factor NDUFAF1 and the OxPhos complex I (CI) core subunit NDUFS3, and plays a critical role in the assembly and stability of CI^25, 26^. ECSIT is also important for the generation of mROS and cellular ROS by macrophages upon detection of pathogens^26^. We have recently generated a mouse strain with floxed *Ecsit*, and demonstrated that loss of ECSIT in macrophages leads to a striking metabolic shift, loss of mitochondrial CI, induction of mitochondrial damage and accumulation of mROS^27^. We have also found that ECSIT interacts with components of the mitochondrial quality control system in macrophages and that ECSIT-deficient cells exhibit decreased mitophagy^27^.

Based on an interaction of ECSIT with presenilin 1 (PSEN1), as well as with redox and mitochondrial proteins encoded by AD susceptibility genes, it was recently proposed that ECSIT could, potentially, integrate amyloid β (Aβ) metabolism, oxidative stress and mitochondrial dysfunction in AD pathogenesis^28, 29^. We hypothesized that dysregulation of ECSIT in neurons might contribute to the pathogenesis of sporadic AD through the accumulation of mROS and resultant neuropathological oxidative stress. Therefore, we investigated the role of ECSIT in AD development by conditionally deleting ECSIT in neurons. We found that targeted loss of ECSIT in the regions of the brain primarily affected by AD led to CI alterations and mROS accumulation, as well as impaired mitophagy. ECSIT deletion in these neurons led to progressive neuropathology with cognitive decline, synaptic dysfunction, neuronal loss, and indicators of neuropathology including gliosis and tau phosphorylation. Specific reduction of mROS by scavenger expression delayed pathogenesis, implicating mROS from defective mitochondria as a pathogenic factor for the observed neuropathology. Intriguingly, we discovered that enhancing ECSIT expression could improve cognitive function in an amyloid-driven model of AD, while reduced ECSIT expression further exacerbated pathology. ECSIT, therefore, prevents oxidative stress in neurons, the development of neuroinflammation and AD-like pathology. Together, these findings provide strong support for the hypothesis that ECSIT protects against neuropathology by maintaining CI function and mitochondrial quality control. Consistent with this hypothesis, we observed that ECSIT is downregulated in AD-affected brains.

## Results

### Loss of ECSIT in neurons leads to oxidative stress and impaired maturation

We began our studies by examining the expression of ECSIT protein in the brain using an anti-ECSIT antibody^23, 27^. ECSIT is expressed broadly throughout the adult mouse cortex and is highly expressed in the pyramidal layer of the hippocampus and the amygdala (Fig. 1a and Extended Data Fig. 1a). As expected, ECSIT is enriched in mitochondria of cortical and hippocampal cells (Extended Data Fig. 1b). Western blotting and immunofluorescence demonstrated that ECSIT is expressed in primary neurons, astrocytes and microglia and exhibits mitochondrial localization (Extended Data Fig. 1c-e). Thus, ECSIT is primarily a mitochondrial protein in neurons, consistent with observations made in other cell types^25, 26^.

**Fig. 1.**
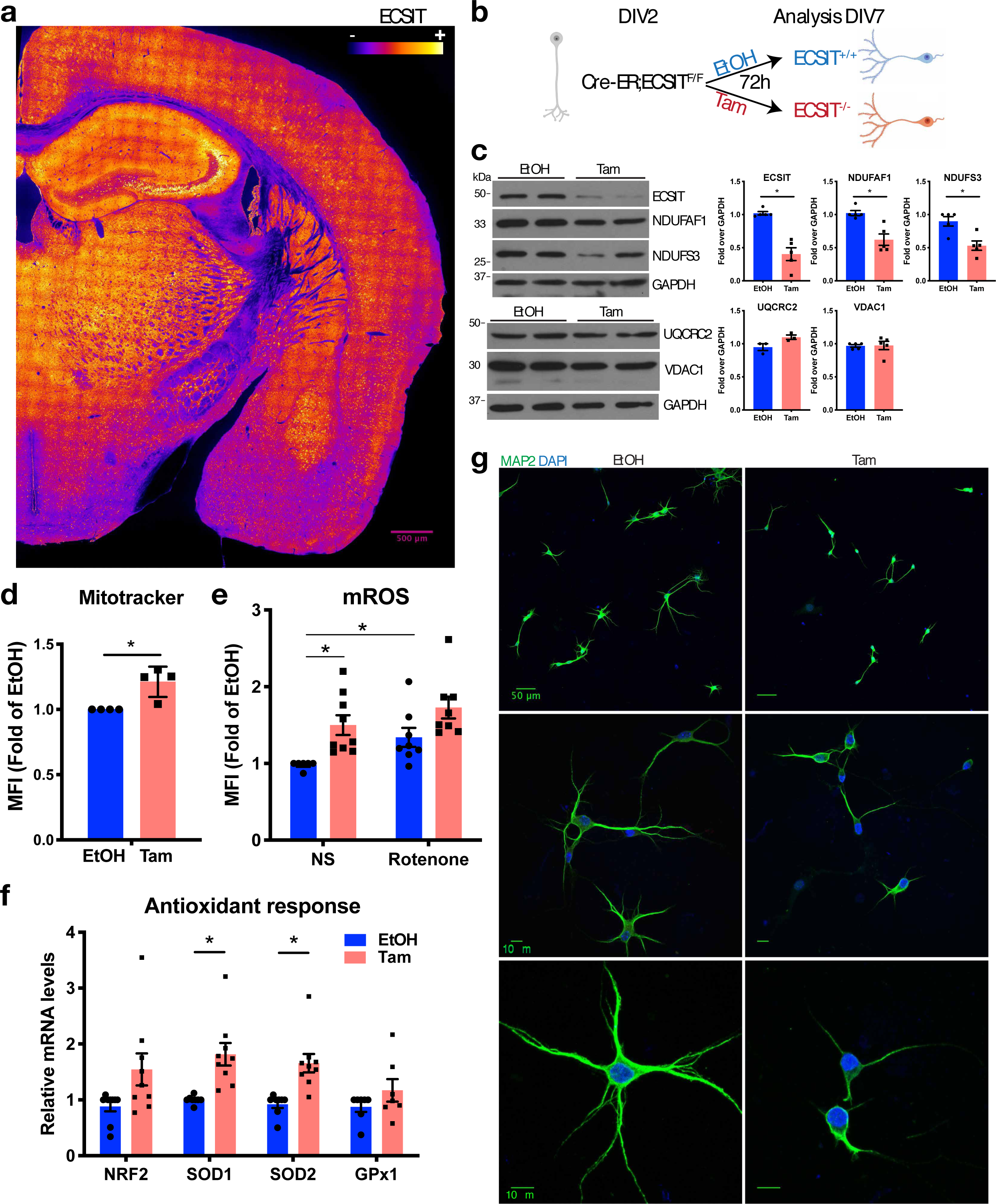
Loss of ECSIT in neurons leads to oxidative stress. **a,** Heatmap showing relative intensity of ECSIT immunostaining in coronal section from an 8-month-old animal. Image processed with Fiji and Fire look-up table (LUT). **b,** Deletion of ECSIT in primary neurons, with indicated genotype, during maturation by activation of Cre with Tamoxifen (Tam) treatment *in vitro*, vehicle: EtOH. **c,** Levels of ECSIT, complex I (NDUFS3, NDUFAF1), complex III (UQCRC2) and mitochondrial porin (VDAC1) assessed by Western blotting of 2 lysates per condition from primary neurons maintained in culture for 7 days *in vitro* (DIV7) and treated as in **b**. Representative of 3 independent experiments, quantified on the right. *, p<0.05 in Mann-Whitney test. **d,** Mean fluorescence intensity (MFI) of mitochondrial marker Mitotracker Green in DIV7 primary neurons treated as in **b** and analyzed by flow cytometry. Mean+/-sem of fold MFI over EtOH of 4 independent experiments. *, p<0.05 in Mann-Whitney test. **e,** Fold MFI of mitochondrial ROS probe MitoSOX over EtOH analyzed by flow cytometry (NS: non-stimulated, rotenone: inducer of ROS from Complex I, positive control). Mean+/-sem of fold MFI over EtOH of 6 experiments is shown. *, p<0.05 in Kruskal-Wallis test with Dunn’s post hoc test. **f,** mRNA level of antioxidant response genes by qPCR in DIV7 primary neurons. NRF2 is a transcriptional master regulator. SOD2 and Gpx1 are located in mitochondria. Average of 8 independent experiments; *, p<0.05 in multiple t-test. **g,** MAP2 and DAPI staining of primary neurons at DIV7. Representative of 5 independent experiments. Scale bar indicated for each pair of images.

To assess the function of neuronal ECSIT, we prepared primary cultures of hippocampal neurons isolated from Cre-ER\**Ecsit^F/F^* neonates. Treatment with tamoxifen (Tam) to induce Cre-mediated *Ecsit* deletion (Fig. 1b) led to decreased ECSIT expression and a concomitant decrease in CI protein levels, compared to vehicle-treated (ethanol; EtOH or Et) primary neurons (Fig. 1c). The decrease in CI proteins was not due to a loss of mitochondria but instead was accompanied by an increase in mitochondrial mass (Fig. 1d and Extended Data Fig. 1i). Levels of mRNA encoding enzymes involved in OxPhos (*Sdha*, which encodes a subunit of complex II, and *Atp5a1*, which encodes a subunit of complex V) and glycolysis (*Pkm2*, *Pfkfb3*, *Mct4*, *Glut1*) were altered, suggesting a shift towards increased glycolysis (Extended Data Fig. 1g). Deletion of *Ecsit* further resulted in increased mROS production, both at steady state (NS) and upon induction of mROS production by rotenone (Fig. 1e and Extended Data Fig. 1f). This was concomitant with an activation of the antioxidant response, reflected in increased transcription of the genes encoding transcription factor NRF2 and the detoxifying enzymes cytosolic SOD1, mitochondrial SOD2 and GPx1 (Fig. 1f). These results indicate that loss of ECSIT in neurons leads to an accumulation of dysfunctional mitochondria and oxidative stress, both of which are detrimental to neuronal function^30^. Deletion of ECSIT also impaired maturation of primary neurons *in vitro*, as quantified by Sholl analysis to evaluate neuronal morphology (Fig. 1g and Extended Data Fig. 1h)^31^.

### Deletion of ECSIT in neurons leads to mitochondrial dysfunction and oxidative stress *in vivo*

To address the role of ECSIT in brain function, we crossed *Ecsit^F/F^* animals with a CamKII-Cre mouse strain. This resulted in deletion of *Ecsit* in excitatory neurons in the hippocampal CA1 pyramidal layer starting at 2.5 weeks, and subsequently in the forebrain^32, 33^. As expected, reduced ECSIT levels were evident in cortex, hippocampus and amygdala and a concomitant decrease of CI proteins (NDUFS3, NDUFAF1), but not of the mitochondrial porin VDAC1, was observed (Fig. 2a and Extended Data Fig. 2a). These *CamKII-Cre*Ecsit^F/F^* animals (CamKII KO) are viable but die prematurely at a median age of 8 months (Extended Data Fig. 2b). Consistent with a defect in CI, neuronal mitochondria in permeabilized synaptosomes isolated from forebrains of CamKII KO mice had decreased mitochondrial potential in the presence of CI substrates, comparable to mitochondria uncoupled by treatment with carbonyl cyanide *m*-chlorophenyl hydrazine (CCCP) (Fig. 2b)^34^. Thus, deletion of ECSIT leads to CI dysfunction and failure to maintain mitochondrial potential in neurons *in vivo*. Moreover, mitochondria accumulated in CamKII KO brain regions where *Ecsi t* had been deleted: namely cortex and amygdala, with a similar trend in hippocampus, but not in midbrain. This was assessed by the ratio of mitochondrial DNA to nuclear DNA (Fig. 2c), and confirmed by the levels of mitochondrial proteins VDAC1, TOM20 and HSP60 in cortex (Extended Data Fig. 2c), demonstrating that loss of CI function is not due to loss of mitochondria. CI subunits were not decreased at the mRNA level (Extended Data Fig. 2d), indicating that NDUFAF1 and NDUFS3 are lost through a posttranslational mechanism, consistent with observations in macrophages^27^.

**Fig. 2.**
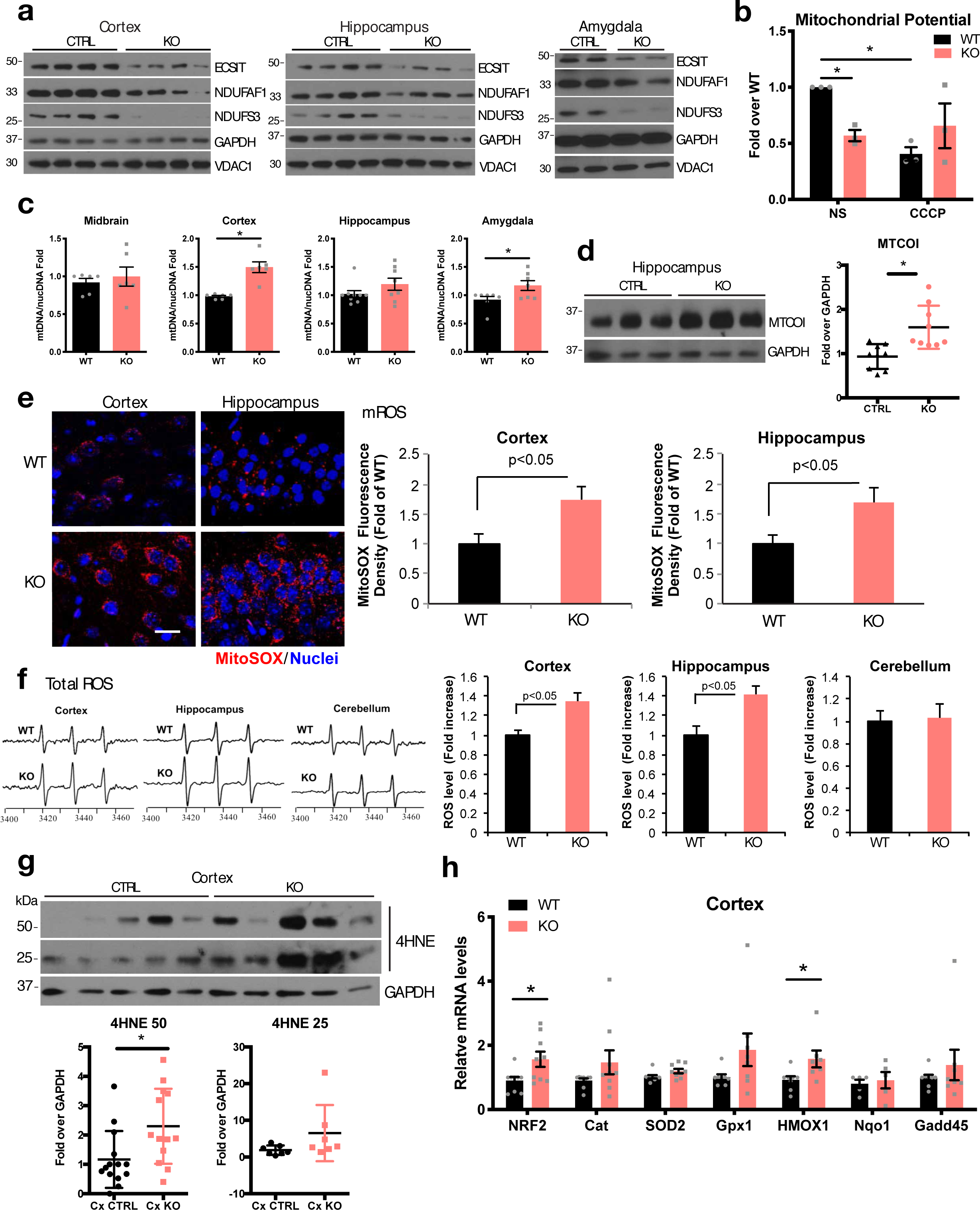
Deletion of ECSIT in neurons *in vivo* leads to mitochondrial dysfunction and oxidative stress in the brain. Animal genotypes: KO=CamKII-Cre^+^ ECSIT F/F or F/-; CTRL=CamKII-Cre^+^ ECSIT+/+ or +/-, and WT=CamKII-Cre^+^ ECSIT+/+. All experiments were performed with 3- to 5-month-old mice. **a,** Western blot analysis of the levels of ECSIT, Complex I (NDUFS3, NDUFAF1) and mitochondrial porin (VDAC1), in lysates from indicated brain regions. Each lane represents an animal. Representative of 3 independent experiments. **b,** Mitochondrial potential levels assessed by TMRM staining and fluorescence reading in permeabilized synaptosomes isolated from cortex+hippocampus homogenates from WT and KO animals, in the presence of complex I substrates (Glutamate+Malate) with and without CCCP treatment. Data are expressed as fold-increase relative to WT. Average of n=3 experiments; *, p<0.05 in one sample t-test. **c,** Mitochondrial mass assessed by the ratio of mitochondrial genomes (*mtCOI* gene) copy numbers over nuclear genomes (*Ndufv1* gene) by qPCR in indicated brain regions. Data are expressed as fold-increase relative to WT. Average of 6 mice per group; *, p<0.05 in Mann-Whitney test. **d,** Western blot analysis for Complex IV subunit MTCOI levels in hippocampal lysates. Quantification of band intensity over GAPDH on the right. Average of 9 mice per group; *, p<0.05 in Mann-Whitney test. **e,** Mitochondrial ROS. (Left) representative image of MitoSOX signals in acute brain slices after *in vivo* staining; scale bar = 25 µm. (Right) quantification of intensity of MitoSOX staining in the indicated regions. Data are expressed as fold increase relative to WT mice. N=3 mice per group. **f,** ROS in different brain regions of WT and KO animals. (Left) representative EPR spectra. (Right) Quantification of EPR spectra in brain homogenates. Data are expressed as fold-increase relative to WT mice. N=3 mice per group. **g,** Representative Western blot analysis of 4-HNE signal revealing peroxidated lipids. Western blot bands are quantified over GAPDH below. Data are expressed as fold increase relative to WT. N=13 (50 kDa) and n=7 (25 kDa) mice per group; *, p<0.05 in Mann-Whitney test. **h,** qPCR levels of antioxidant response genes in cortex lysates. Average of n=8 mice per group, *, p<0.05 in multiple t-test.

Increases in the levels of other respiratory subunits suggested a compensatory upregulation of other complexes, such as complex IV represented by the MTCOI protein (Fig. 2d and Extended Data Fig. 2d). The effects of ECSIT loss were more evident in cortex and amygdala than in hippocampus. However, it should be noted that *Ecsit* deletion by CamKII-Cre is mainly targeted to the CA1 region of the hippocampus (Fig. 3e)^32^, while it is expressed at higher levels in CA2-3 regions (Extended Data Fig. 1a). Maintenance of ECSIT and CI levels in CA2-CA3 regions may obscure the effects of *Ecsit* deletion in these assays, which use the entire hippocampus. Thus, both *in vitro* and *in vivo*, targeted, neuronal deletion of *Ecsit* caused impairment of CI function and simultaneous accumulation of mitochondria.

**Fig. 3.**
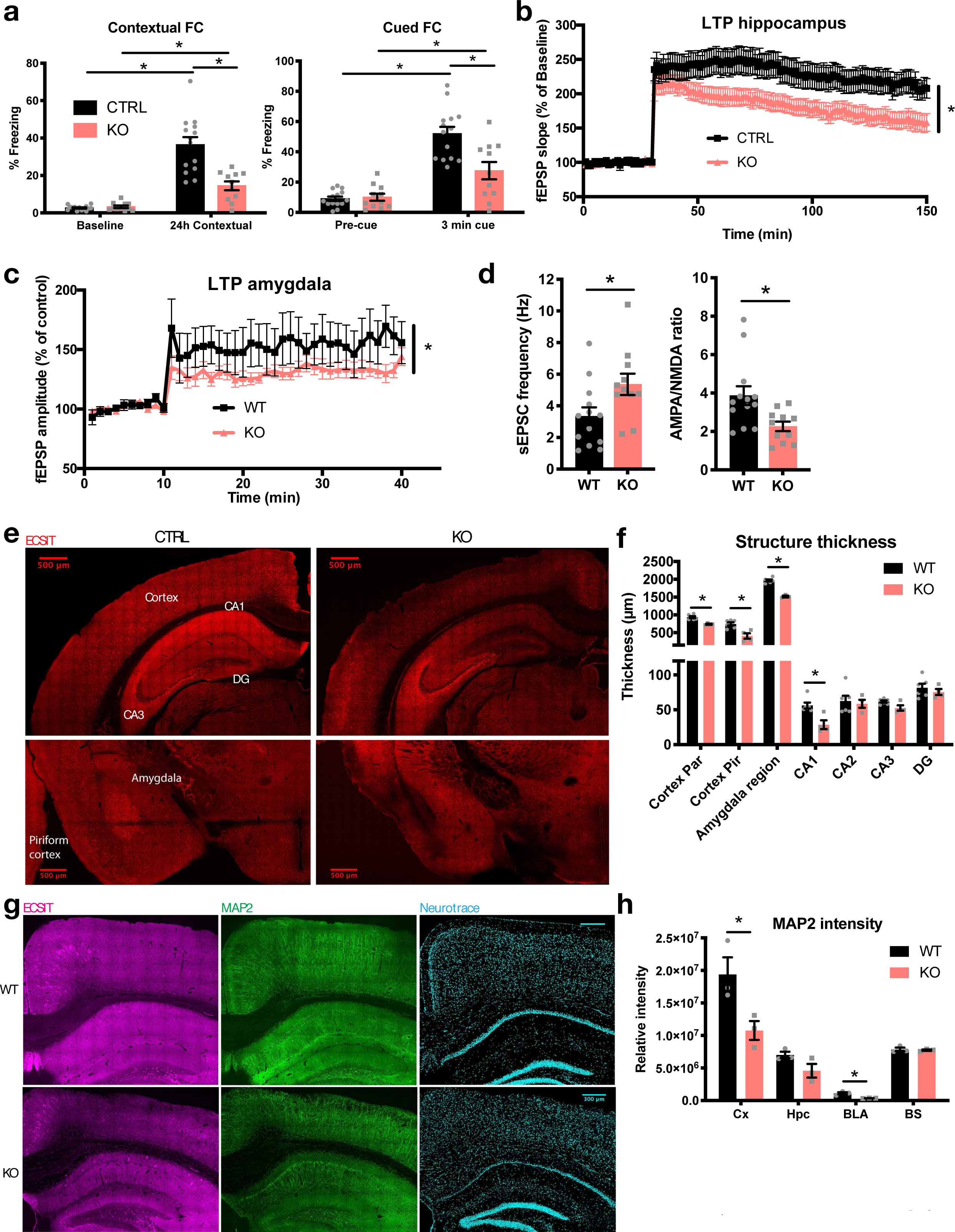
ECSIT deletion in neurons leads to cognitive dysfunction and neurodegeneration. **a,** Contextual and cued fear conditioning (FC) in 3- to 5-month-old animals. Freezing behavior was evaluated in response to context first (24h contextual), then to cue (3 min cue). n=14 CTRL;14 KO, *, p<0.05 in 2-way ANOVA with Sidak’s post-hoc test. **b,** Long-term potentiation (LTP) in hippocampal acute slices from 3- to 5-month-old animals (n=17 CTRL; 21 KO slices), *, p<0.05 in 2-way ANOVA with repeated measures after tetanus. **c,** Electrophysiology in basolateral amygdala (BLA) in 3- to 5- month-old animals (n=13 WT; 11 KO slices). *, p<0.05 in 2-way ANOVA with repeated measures after tetanus. **d,** Basal synaptic transmission in BLA as indicated in the same samples as in **c**. *, p<0.05 in t-test. **e,** Immunostaining for ECSIT and confocal microscopy in coronal sections of 8-month-old animals. Representative of 6 animals per group. **f,** Quantification of structure thickness from Neurotrace staining of coronal brain sections (Par=parietal, Pir=piriform, DG=dentate gyrus) of 8- to 9-month-old animals. N=6 WT and 4 KO. *, p<0.05 in t-test. **g,** Representative immunofluorescence staining of MAP2 and confocal acquisition in hippocampus and cortex. **h,** Quantification of MAP2 intensity of three 8-month-old animals per group (Cx=Cortex, Hpc=Hippocampus, BS=brainstem).

*Ecsit* deletion leads to the accumulation of mROS as well as increased oxidative stress in primary neurons (Fig. 1e-f) in agreement with observations in immortalized cells^27^. Similarly, H_2_O_2_ ROS measured with Amplex Red in isolated mitochondria in the presence of CI substrates was higher in KO than in WT condition (Extended Data Figure 2e). Consistently, *in vivo* evaluation of mROS using MitoSOX demonstrated increased mROS in the cortex and hippocampus of CamKII KO animals compared to controls (Fig. 2e). This was accompanied by an increase in total ROS in the regions with *Ecsit* deletion, but not in areas with unaltered ECSIT expression, such as the cerebellum (Fig. 2f). ROS can induce lipid peroxidation (LPO), generation of the reactive lipid 4-hydroxynonenal (4-HNE) and increased 4-HNE adducts. Increased levels of 4-HNE therefore are considered evidence of oxidative stress in AD brains^35^.

Consistent with direct ROS measurements, we observed increased levels of LPO in the cortex of CamKII KO animals (Fig. 2g). The accumulation of ROS led to the induction of an antioxidant response in cortex, amygdala and hippocampus, as evidenced by qPCR analysis of the oxidative stress response master regulator NRF2 and several antioxidant enzymes (Fig. 2h and Extended Data Fig. 2f). These results confirm that *Ecsit* deletion in neurons leads to oxidative stress in the CNS.

### *Ecsit* deletion induces hippocampal and amygdala electrophysiological dysfunction

Since excessive oxidative stress may perturb neuronal processes^30^, we hypothesized that *Ecsit* deletion in hippocampal and amygdala neurons would lead to progressive cognitive dysfunction. We therefore investigated the ability of CamKII KO mice to perform in tasks reliant on hippocampal and amygdala function. Evaluation of associative memory by the fear conditioning (FC) test^36^ revealed a memory deficit in CamKII KO mice compared to WT controls during both contextual and cued fear learning assessments (Fig. 3a). These defects were not accompanied by deficits in the sensory threshold that might have affected the capability of the mice to perceive electric shock (data not shown). Spatial memory evaluation by the 2-day Radial Arm Water Maze test (2-day RAWM) revealed no difference between controls and CamKII KO mice (Extended Data Fig. 3a).

The FC learning paradigm depends on the hippocampus and amygdala. The hippocampus, in particular, is indispensable for contextual FC, whereas the amygdala is involved both in contextual and cued memory^37^. These findings prompted us to analyze long-term potentiation (LTP), an electrophysiological correlate of memory, both in hippocampus and amygdala. Indeed, we observed reduced LTP at CA1-CA3 synapses (Fig. 3b). Basal transmission, however, was normal in the hippocampus, as assessed by measurement of input/output and paired-pulse facilitation (PPF), a form of short-term plasticity (Extended Data Fig. 3b). In the basolateral amygdala (BLA), LTP was also reduced (Fig. 3c) and basal synaptic transmission was affected (Fig. 3d). Older animals (6- to 8-month-old) exhibited an exacerbated phenotype: the deficits in cognitive function during FC were accompanied by deficits in 2-day RAWM performance and defects in visual-motor-motivational parameters in the visible platform task (Extended Data Fig. 3c-e). Furthermore, synaptic function of the hippocampus was inhibited during both LTP and basal synaptic transmission (Extended Data Fig. 3f-g). As CamKII promoter-driven Cre expression spreads to the forebrain, and has reached maximum levels by 2 months of age^32, 33^, these findings do not reflect progressive deletion of *Ecsit* over time but rather indicate a progressive decline in cognitive and electrophysiological function resulting directly from loss of neuronal *Ecsit* expression.

### *Ecsit* deletion in excitatory neurons leads to neuropathology and neuroinflammation

To determine whether this progressive cognitive impairment also manifested as changes to the brain structure, we examined brain sections of 8-month-old animals by immunofluorescence. We observed a significant thinning of the CA1 layer of hippocampus and the parietal section of cortex, as well as atrophy of the BLA (Fig. 3e-f and Extended Data Fig. 3h). The organization of cortical layers was also disrupted (Extended Data Fig. 3h, j). Thus, all regions to which *Ecsit* deletion was targeted were affected, while regions without deletion, i.e. CA2, CA3 and the dentate gyrus, were spared (Fig. 3e-f and Extended Data Fig. 3h, j). Moreover, MAP2 expression, a marker of neurites, was decreased in the cortex, CA1 neuronal dendrites and BLA (Fig. 3g-h and Extended Data Fig. 3i). These phenotypes were not observed in younger, 3- to 5-month-old animals (data not shown), consistent with the progressive nature of the neuropathology and brain atrophy caused by targeted deletion of *Ecsit* in neurons.

A hallmark of neurodegenerative diseases is the development of neuroinflammation, which is linked to gliosis, i.e. the accumulation of astrocytes and microglia^38^. Using immunofluorescence, we observed increased staining for glial fibrillary acidic protein (GFAP) and ionized calcium binding adapter molecule (Iba1) in the cortex and hippocampus from 8-month-old CamKII KO mice, suggesting accumulation of astrocytes and microglia respectively (Fig. 4a). mRNA levels of neuroinflammatory markers (CD68 for activated microglia, as well as the proinflammatory cytokines TNF, IL-1α, and IL-1β) were also increased in the hippocampus of CamKII KO animals (Fig. 4b). Phosphorylation of tau protein at multiple residues, which is thought to regulate tau microtubule binding activity, aggregation, subcellular distribution and signaling functions^39^, has been demonstrated to be increased in AD brains^40^. Using antibodies against discrete AD-related phospho-tau epitopes (PHF1: S396/S404 and CP13: pS202), we observed increased signals in hippocampal and cortical lysates from CamKII KO animals, with no significant change in total soluble tau (Fig. 4c). These results are consistent with neuronal damage and AD-related neuropathological processes.

**Fig. 4.**
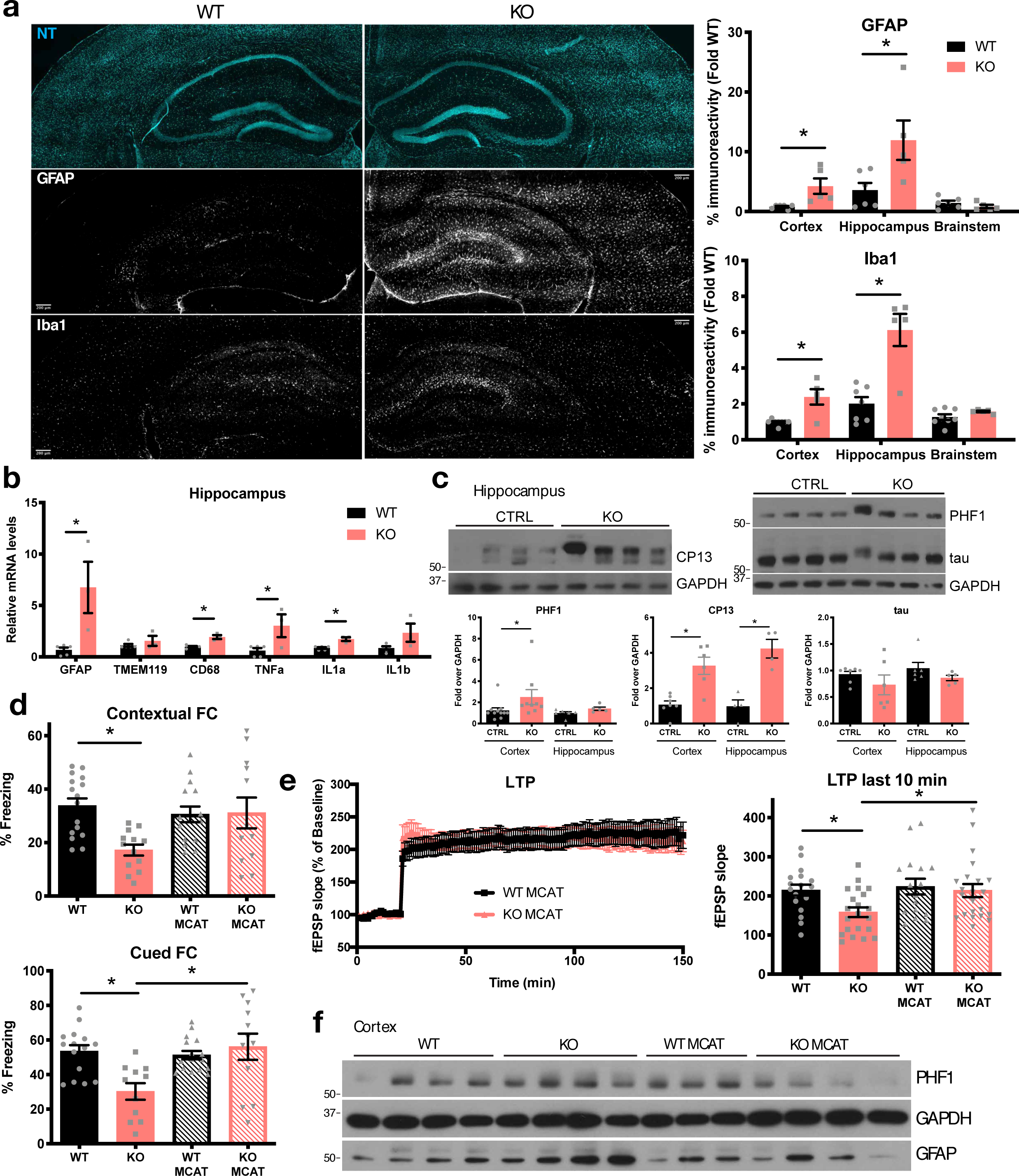
Effects of ECSIT deletion in the CNS are rescued by scavenging mROS. **a,** Gliosis analysis by immunofluorescence staining of GFAP (astrocytes) and Iba1 (microglia) and confocal acquisition in coronal sections of brains from 8-month-old animals. NT, neurotrace. (Right) quantification of percent immunoreactivity of indicated staining and areas. N=5 WT and 3 KO. *, p<0.05 in Mann-Whitney test in each region. **b,** mRNA levels of indicated genes by qPCR in hippocampus of 5- to 6-month-old animals. N=5 WT; 3 KO. *, p<0.05 in Mann-Whitney test. **c,** Western blot analysis of total soluble tau and phosphorylated tau epitopes PHF1 and CP13 in hippocampus lysates from 3- to 5-month-old animals. Quantification of band intensities in cortex and hippocampus lysates over GAPDH (n=4-9 animals per groups as indicated). *, p<0.05 in Mann-Whitney test in each region. **d,** Fear conditioning (FC) in MCAT-expressing animals. Baseline in Extended Data Figure 4g. N=16 WT; 13 KO; 15 WT MCAT (WT with MCAT transgene); 12 KO MCAT (KO with MCAT transgene). *, p<0.05 in Kruskal-Wallis test with Dunn’s post hoc test. **e,** LTP in hippocampus from 3- to 5-month-old animals. Comparison of the last 10 min of LTP between the 4 groups (n=17; 21; 16; 22, resp.). *, p<0.05 in Kruskal-Wallis test with Dunn’s post hoc test. **f,** Levels of phosphorylated tau epitope PHF1 and GFAP assessed by Western blot of cortical lysates from 3- to 5-month-old animals.

### ECSIT protects the brain from mROS-induced neuropathology

To conclusively demonstrate causal involvement of mROS in neuropathology following *Ecsit* deletion, we attempted to rescue the phenotype of CamKII KO mice by scavenging mROS using the MCAT transgenic mouse model^41^. In this mouse strain, human catalase is overexpressed in mitochondria, leading to potent elimination of mitochondrial hydrogen peroxide. We observed reliable expression of the MCAT transgene in all brain regions (Extended Data Fig. 4a). Remarkably, cognitive dysfunction, assessed by fear conditioning, was rescued in CamKII KO animals expressing MCAT (KO MCAT, Fig. 4d and Extended Data Fig. 4b). Moreover, LTP in hippocampus was also restored: there was no difference in the amount of potentiation between WT, WT MCAT and KO MCAT animals (Fig. 4e). Finally, scavenging mROS by MCAT overexpression curbed tau phosphorylation and gliosis in CamKII KO mice (Fig. 4f and Extended Data Fig. 4c). These data clearly establish mROS as a major mediator of the phenotypic changes and neuropathology resulting from decreased neuronal ECSIT expression.

We prepared primary neurons from *Cre-ER*Ecsit^F/F^* mice either expressing MCAT or not, and inducibly deleted *Ecsit* by tamoxifen (Tam) treatment, as in Figure 1. As expected, MCAT expression prevented the increase in mROS in *Ecsit* KO neurons, as measured by MitoSOX staining in *Ecsit* KO MCAT (Tam MCAT) neurons compared to WT MCAT (Et MCAT) neurons after 7 days of culture *in vitro* (Extended Data Fig. 4d). However, scavenging mROS did not rescue the defect in *in vitro* maturation of primary neurons (Extended Data Fig. 4e), suggesting an additional role for ECSIT in neuronal function beyond mROS control. We have previously shown that ECSIT is critical for mitophagy in macrophages^27^. In primary neurons and in brains from mice with deletion of *Ecsit* in neurons we observed that mitochondria accumulated despite mitochondrial dysfunction, indicating impaired mitophagy (Fig. 1 and 2). Furthermore, MCAT expression did not rescue mitochondria accumulation in the cortex of CamKII KO mice (Extended Data Fig. 4f). This prompted us to ask whether ECSIT was also involved in neuronal mitophagy.

### Loss of neuronal ECSIT disrupts mitophagy

Mitophagy describes the selective removal of dysfunctional mitochondria by autophagy, and failure to eliminate damaged mitochondria is believed to contribute to increased levels of mROS. Recruitment of the autophagosomal protein LC3b to mitochondria is a key indicator of mitophagy^43^, so we analyzed LC3b localization by immunofluorescence after *Ecsit* deletion. We observed a striking difference in LC3b distribution in the soma of primary neurons, with a diffuse pattern in vehicle-treated control neurons (EtOH) and the presence of aggregates in *Ecsit* KO cells (Tam) (Fig. 5a, top). These aggregates in the soma did not colocalize with mitochondria, which were identified by staining for the mitochondrial outer membrane protein TOM20. In neuronal processes, however, a majority of the mitochondria were associated with clustered LC3b in *Ecsit* KO neurons, but not in control neurons (Fig. 5a, bottom, pink arrows, and Extended Data Fig. 5a). This suggested recruitment of the autophagy machinery and mitophagy induction without completion of autophagic degradation. Mitophagy can be inhibited by increased mitochondrial size resulting from defective mitochondrial fission, oxidative stress or metabolic adaptations^44, 45^. However, quantification of mitochondrial length in neuronal processes *in vitro* showed that mitochondria of *Ecsit* KO neurons were shorter, rather than longer, than the mitochondria of control neurons (Extended Data Fig. 5b).

**Fig. 5.**
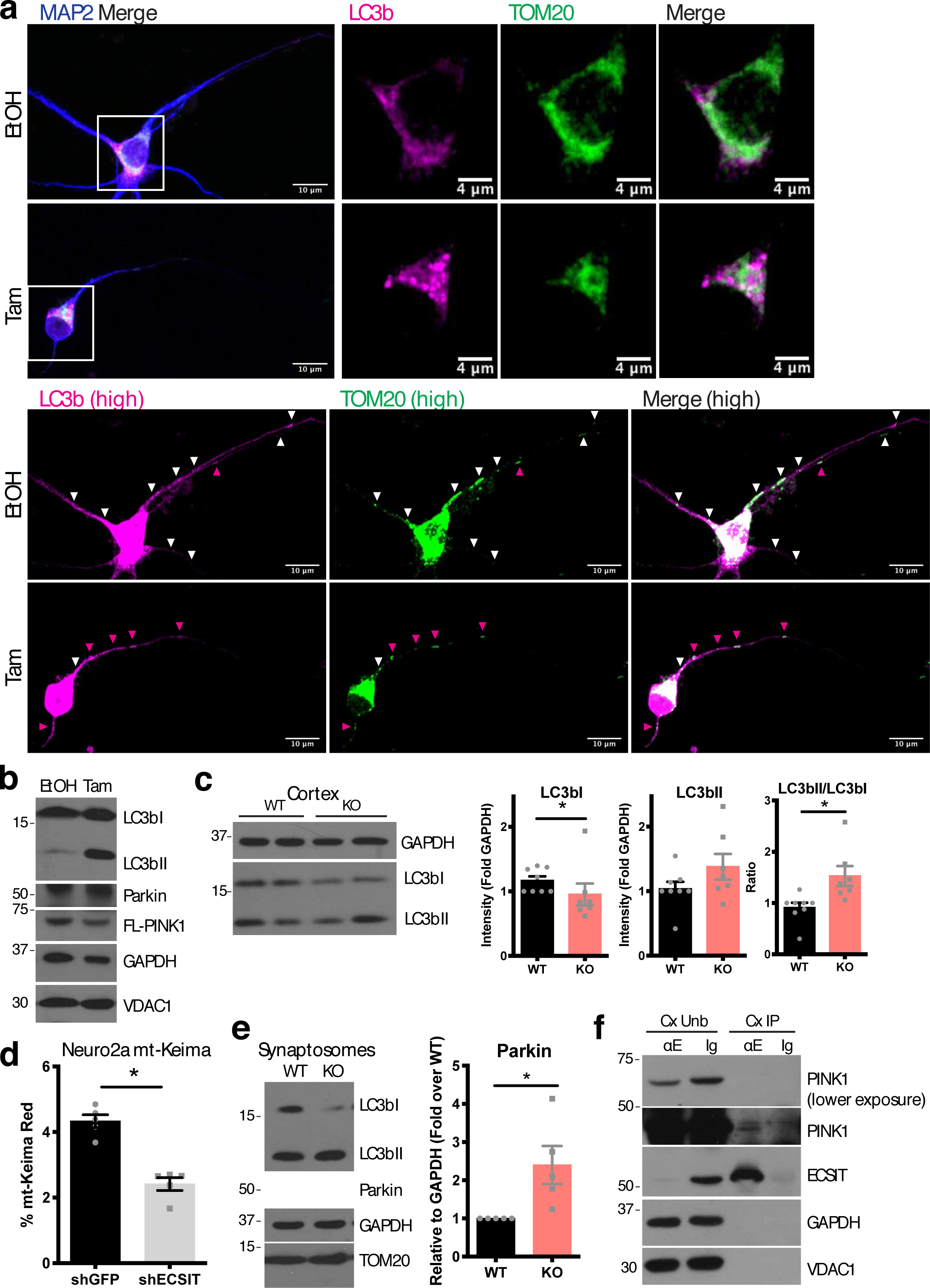
Loss of ECSIT in neurons leads to mitophagy alteration. **a,** MAP2, LC3b and TOM20 immunofluorescence staining and confocal microscopy in DIV7 primary neurons as in Fig. 1b. Images were processed with Fiji to enhance signal (high) and merge channels (white signal). Magenta arrows indicate mitochondria with enriched LC3b as opposed to white arrows, mitochondria with no LC3b enrichment. Representative images of at least 12 cells per condition in 3 independent experiments, quantified in Extended Data Fig 5a. **b,** Representative Western blot analysis of LC3b, Parkin and full-length (FL-)PINK1 levels in DIV7 primary neurons (Fig. 1b) of 3 independent experiments. Et, ethanol; Tam, Tamoxifen. **c,** Western blot analysis of LC3bI and LC3bII levels in cortical lysates from 3- to 5-month-old mice. (Left) representative experiment. (Right) quantification of intensities from n=8 WT;7 KO. *, p<0.05 in Mann-Whitney test. **d,** Percent of mt-Keima “Red” cells in Neuro2A control (shGFP) or ECSIT downregulated by shRNA (shECSIT) stably expressing the mt-Keima mitophagy reporter, analyzed by flow cytometry. Average of 5 independent experiments. *, p<0.05 in t-test. **e,** Western blot analysis of Parkin and LC3b levels in synaptosomal mitochondria isolated from brain homogenates from 3- to 5-month-old animals. (Left) representative experiment. (Right) quantification of Parkin band intensities over GAPDH for 5 animals per group. *, p<0.05 in Mann-Whitney test. **f,** Coimmunoprecipitation of PINK1 with ECSIT assessed by Western blot in adult WT mouse cortex (Cx) lysates. Unb, unbound: cell lysate supernatant after immunoprecipitation with the indicated antibodies; IP, fraction immunoprecipitated with anti-ECSIT antibody (E) or IgG control (Ig). Representative of 3 independent experiments. The same experiment is represented in Extended Data Fig. 6h.

Upregulation of ATG7, an early regulator of autophagy, and LC3b mRNA levels in *Ecsit* KO neurons confirmed that general autophagy induction was not inhibited and indicated a cellular response to mitochondrial dysfunction (Extended Data Fig. 5c). LC3b can be used as a marker of progression through autophagy, as it undergoes modification from LC3bI to lipidated LC3bII and finally lysosomal degradation after autophagy activation^43^. In agreement with the immunofluorescence results, we observed accumulation of LC3bII in primary *Ecsit* KO neurons (Fig. 5b). Similarly, we observed an accumulation of LC3bII relative to LC3bI in the cortex of CamKII KO mice (Fig. 5c), suggesting that the autophagic flux is altered in CamKII KO brains and results in mitophagy defects.

The mitophagy reporter mt-Keima is a ratiometric indicator of mitophagy completion, which exhibits red fluorescence in the acidic environment of the lysosome^46^. Using shRNA, we downregulated ECSIT in mt-Keima-expressing Neuro2a cells. ECSIT knockdown (shECSIT) replicated the mitochondrial accumulation (VDAC1 and TOM20) and LC3bII accumulation observed in ECSIT KO primary neurons (Extended Data Fig. 5d). We observed a decrease in mt-Keima red fluorescence (Fig. 5d), confirming a defect in mitophagy completion. The best-understood mitophagy pathway is triggered by acute mitochondrial damage and involves stabilization of full-length (FL)-PINK1 at the outer membrane of damaged mitochondria, accumulation of the ubiquitin ligase Parkin, ubiquitination of mitochondrial targets and recruitment of LC3bII and the autophagy machinery^47, 48^. We have previously shown that ECSIT is ubiquitinated upon induction of mitochondrial depolarization by CCCP in macrophages and is a substrate for ubiquitination by Parkin^27^. Although mitophagy can be acutely triggered by mitochondrial damage, constant basal mitophagy to enable mitochondrial turnover has also been observed in macrophages^27^ and brain tissues^12, 22, 46, 49, 50^. In primary neurons, we found that *Ecsit* deletion caused upregulation of Parkin (Fig. 5b and Extended Data Fig. 5c), similar to our observations in macrophages^27^. Synaptosomes are enriched for neuronal mitochondria^51^, and we observed increased Parkin levels in synaptosomes isolated from the forebrains of CamKII KO mice compared to those from WT mice (Fig. 5e). This demonstrates that the induction of mitophagy in response to mitochondrial damage remains intact in the absence of ECSIT. As in macrophages, we found that ECSIT interacts with FL-PINK1 at steady-state in cortical lysates from WT mice (Fig. 5f)^27^. Interestingly, although mRNA levels of PINK1 were not significantly altered in primary neurons deleted for ECSIT, there was a trend towards decreased FL-PINK1 protein levels (Fig. 5b). Similarly, we observed decreased FL-PINK1 in the cortex from CamKII KO animals (Extended Data Fig. 5e), consistent with observations in hippocampal tissue from AD brains^22^. Thus, loss of ECSIT in neurons causes an altered autophagic flux, defective mitophagy and accumulation of damaged mitochondria.

### Downregulation of ECSIT expression in the context of AD contributes to disease

Our results establish that loss of neuronal ECSIT leads to mitochondrial dysfunction, mROS accumulation, defective mitophagy, neuroinflammation and neuropathology. All of these events have been implicated in AD development^22, 48^, which led us to hypothesize that ECSIT dysregulation could promote AD pathology. We performed Western blot analysis of post-mortem brain tissues from AD patients obtained through two brain banks: 5 sample pairs from Columbia University Alzheimer’s Disease Research Center (ADRC) and New York Brain Bank (NYBB), uniquely diagnosed by local pathological marks; and 12 sample pairs from the Arizona Study of Aging and Neurodegenerative Disorders (AZSAND) and the Banner Sun Health Brain and Body Donation Program (BBDP) (www.brainandbodydonationprogram.org), renowned for ensuring short post-mortem intervals (table S1)^52^. Cortical lysates from AD patient samples showed a significant reduction of ECSIT protein levels compared to matched controls, as well as a consistent reduction of the ECSIT partner NDUFAF1 (Fig. 6a and Extended Data Fig. 6a). Levels of cytosolic (Actin) and mitochondrial (HSP60) control proteins were unaffected, establishing that the decrease of ECSIT and NDUFAF1 was not the result of an overall loss of mitochondria. ECSIT downregulation in AD brain samples was also observed at the mRNA level in ADRC samples (Fig. 6b) and in data from the population-based prospective Hisayama study (p=0.0043, data not shown)^53^. Moreover, analysis of published gene expression data from microdissected healthy entorhinal neurons of post-mortem tissue sections from AD brains showed downregulation of ECSIT in AD samples, independently of the expression of interacting partners such as NDUFAF1, NDUFS3 and TRAF6 (Extended Data Fig. 6b)^54^.

**Fig. 6.**
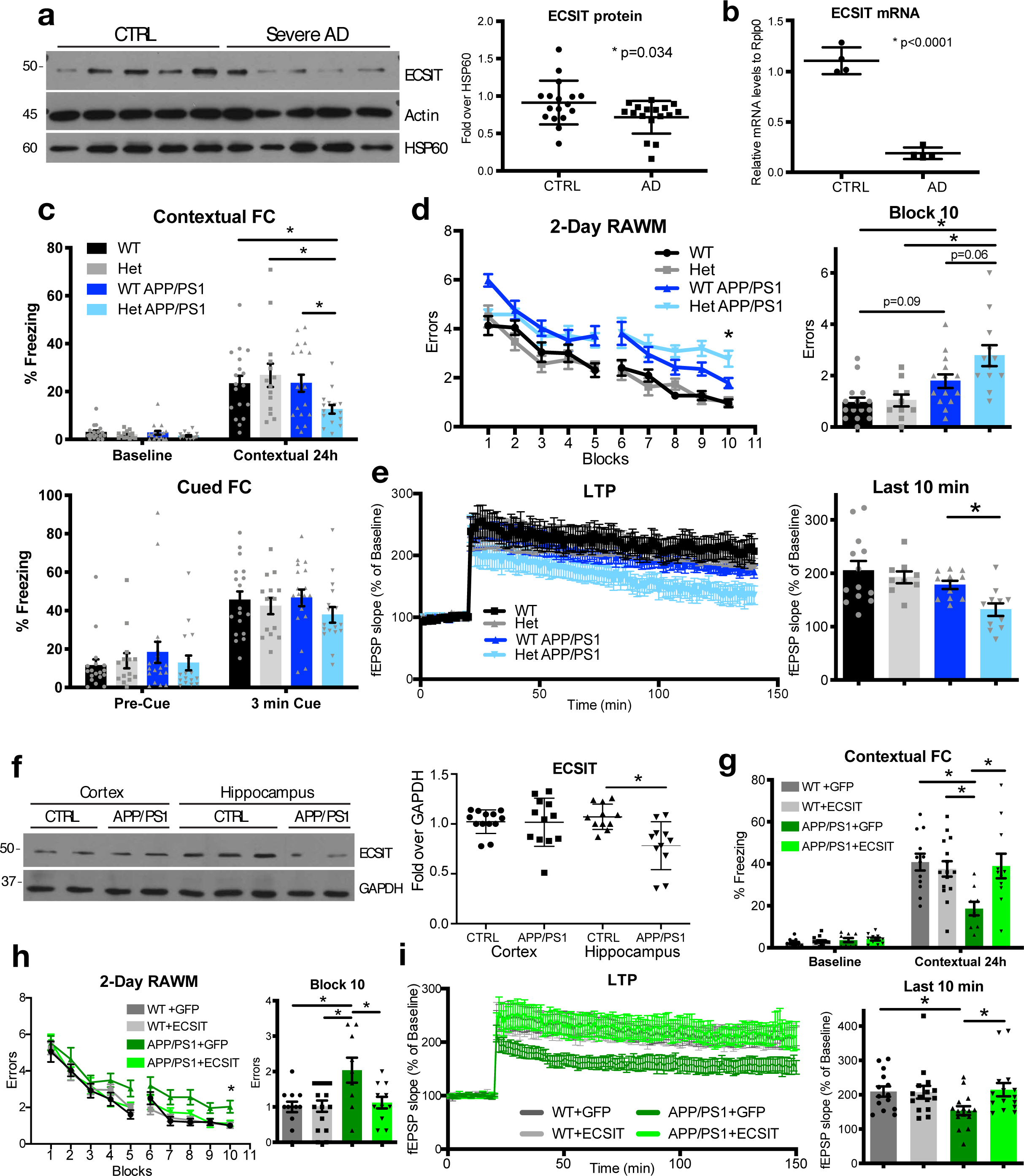
ECSIT downregulation contributes to and ECSIT expression protects against AD-like pathogenesis. **a,** (Left) Representative Western blot analysis of ECSIT levels in human control and AD patient samples and (Right) quantification of band intensities over HSP60 in ADRC and BBDP cohorts. p, p-value in t-test. **b,** mRNA levels of *ECSIT* in ADRC human control and AD patient samples by qPCR. *, p<0.001 in Mann-Whitney test. **c, Animal genotypes: WT**=Synapsin-Cre^+or-^ ECSIT+/+ non-transgenic (Ntg); **ECSIT Het**=Synapsin-Cre^+^ ECSIT F/+ Ntg; **APP/PS1**=Synapsin-Cre^+or-^ ECSIT+/+ with APP/PS1 transgenes; **ECSIT Het APP/PS1**= Synapsin-Cre^+^ ECSIT F/+ with APP/PS1 transgenes. Contextual and cued fear conditioning (FC) on 3- to 5-month-old ECSIT WT or Het, non-transgenic or APP/PS1-transgenic animals. N=18 WT; 14 Het; 17 APP/PS1; 15 Het APP/PS1. *, p<0.05 in ANOVA with Sidak’s post hoc test. **d,** (Left) number of errors per block trial in 2-day RAWM. (Right) statistics for block 10. N=15 WT; 10 Het; 14 APP/PS1; 12 Het APP/PS1 animals. *, p<0.05 in ANOVA with Sidak-Holm’s post hoc test. **e, (Left)** LTP in hippocampus of 3- to 5-month-old animals. (Right) average of the last 10 min for n=13 WT; 9 Het; 11 APP/PS1; 11 Het APP/PS1. *, p<0.05 in Kruskal-Wallis test with Dunn’s post hoc test. **f, (Left)** Levels of ECSIT assessed by Western blot in brain lysates from 2-month-old APP/PS1 animals. (Right) quantification of band intensities over GAPDH for 11 animals per group. *, p<0.05 in Mann-Whitney test for each region. **g,** Contextual fear conditioning (FC) on 5-month-old WT or APP/PS1 animals 3 months after stereotaxic injection of indicated virus. N=12 WT+GFP; 15 WT+ECSIT; 9 APP/PS1+GFP; 11 APP/PS1+ECSIT. *, p<0.05 in 2-way ANOVA with Sidak’s post hoc test. **h,** (Left) number of errors per block of trials in 2-day RAWM; (right) statistics for block 10. Same animals as in **g**. *, p<0.05 in ANOVA with Sidak’s post hoc test. **i,** LTP in hippocampus of 6-month-old animals. (Right) average of the last 10 min. N of acute slices per group=14 WT+GFP; 15 WT+ECSIT; 15 APP/PS1+GFP; 15 APP/PS1+ECSIT. *, p<0.05 in Kruskal Wallis test with Dunn’s post hoc test.

To investigate whether loss of ECSIT can worsen AD-like pathology in the context of Aβ elevation, we used conditional deletion to reduce ECSIT expression in the APP/PS1dE9 amyloid-driven mouse model of AD. To achieve partial downregulation of ECSIT in total brain tissue, as observed in AD patient samples, we crossed heterozygous *Ecsit*-floxed mice with the Synapsin-Cre mouse strain. *Synapsin-Cre*Ecsit^F/+^* (Het) animals are viable and display partially reduced levels of ECSIT in cortex and hippocampus compared to *Synapsin-Cre*Ecsit^+/+^* mice (WT) (Extended Data Fig. 6c). As expected, heterozygous loss of *Ecsit* did not result in behavioral or electrophysiological deficits at 8 months of age (data not shown). We then crossed these mice with the transgenic APP/PS1dE9 mouse strain (abbreviated APP/PS1) bearing the APP KM670/671NL (Swedish) and PSEN1 deltaE9 mutations^55, 56^. *Ecsit* heterozygosity in neurons led to a significant decrease in survival of APP/PS1 transgenic animals (Het APP/PS1) compared to *Ecsit* WT APP/PS1 littermates (WT APP/PS1) (Extended Data Fig. 6d). This finding demonstrates that decreased expression of ECSIT accelerates disease through mitochondrial dysfunction, in parallel to the amyloid-driven neuropathology that follows the overexpression of mutated APP and PSEN1.

Associative memory, tested by contextual FC in mouse models, and spatial memory, tested with 2-day RAWM, are both affected in AD patients^57, 58^. We performed behavioral testing to investigate the cognitive functions of *Ecsit* Het APP/PS1 compared to *Ecsit* WT APP/PS1 mice. We chose an early time point, at which the APP/PS1 mouse model shows only minor cognitive deficits (Fig. 6c-d). At 3 to 5 months of age, we observed that downregulation of ECSIT led to diminished performance in the contextual FC and RAWM tasks for *Ecsit* Het APP/PS1 mice compared to *Ecsit* WT APP/PS1 (Fig. 6c-d). No differences in sensory threshold or visual-motor-motivational parameters were detectable (Extended Data Fig. 6e-f). To investigate the synaptic basis for the memory deficits, we further performed electrophysiological studies in the hippocampus and observed a decrease in LTP (Fig. 6e). Basal synaptic transmission (input/output), but not PPF, exhibited a similar trend (Extended Data Fig. 6g). ECSIT has been reported to interact with PSEN1 in human SH-SY5Y neuronal cells^29^. Analogously, we were able to detect an interaction of ECSIT with PSEN1 in mouse cortical lysates (Extended Data Fig. 6h). However, downregulation of ECSIT had no effect on soluble amyloid β (Aβ_1-42_) levels in APP/PS1 animals (Extended Data Fig. 6i). The APP/PS1 amyloid-driven model of AD does not display some of the pathological marks of AD, such as marked neuronal loss and tangles, which likely require other pathogenic processes^59, 60^. We analyzed whether downregulation of ECSIT triggered the appearance of tau phosphorylation or neuronal loss, as in CamKII KO animals. Immunoblot analysis for tau phosphorylation with the CP13 antibody and neuronal loss with MAP2 antibody did not show significant differences (Extended Data Fig. 6j-k). However, we did observe a trend towards GFAP upregulation, which indicates an increase in astrogliosis. Taken together, these results suggested that there is an additive or synergistic effect of mitochondrial dysfunction, due to decreased ECSIT expression in amyloid-driven AD-like neuropathology.

### ECSIT protects against cognitive decline in AD-like pathogenesis

Our studies show that i) ECSIT is downregulated in AD brain samples; ii) loss of ECSIT leads to neuropathology in mice; and iii) ECSIT downregulation in neurons accelerates cognitive deficits in the APP/PS1 model. These results suggested that preservation of ECSIT expression could have neuroprotective effects. Interestingly, we observed downregulation of ECSIT in hippocampus of 2-month-old APP/PS1 animals (Fig. 6f), consistent with the downregulation of ECSIT mRNA seen in another APP/PS1 model^56^. These early changes in ECSIT mRNA levels were independent of changes in mitochondrial mass (Extended Data Fig. 7a) and expression of other CI proteins (data not shown). This indicates that ECSIT downregulation preceded the development of mitochondrial dysfunction and pathology in this model. Therefore, we investigated whether maintenance of ECSIT expression in the hippocampus would delay or prevent the onset of pathology in the APP/PS1 animals. We used an AAV9 vector driving expression of ECSIT and GFP from a Synapsin promoter to specifically increase ECSIT levels in neurons^61^. Using this approach to infect primary neurons *in vitro* resulted in overexpression of ECSIT with mitochondrial localization, based on colocalization with the mitochondrial protein TOM6 (Extended Data Fig. 7b). We used stereotaxic injection into the brain of 2-month-old mice to augment ECSIT expression in neurons *in vivo*. Three weeks later, GFP and ECSIT expression were analyzed by immunofluorescence in brain sections showing overexpression of ECSIT in the hippocampus (Extended Data Fig. 7c, d). Three months after ECSIT or control GFP AAV injection, APP/PS1 animals were analyzed for cognitive function. As expected, APP/PS1 animals injected with GFP AAV control showed defects in contextual FC (Fig. 6g). Remarkably, ECSIT overexpression rescued this defect. Cued FC was not changed at that age (Extended Data Fig. 7e). ECSIT overexpression also ameliorated spatial memory deficits of APP/PS1 animals in the 2-day RAWM assay (Fig. 6h). Of note, ECSIT overexpression did not modify cognitive function in these behavioral tasks in WT animals (WT+ECSIT compared to WT+GFP) (Fig. 6g-h), and sensory-motor and motivational controls did not exhibit significant differences between any of the groups (Extended Data Fig. 7f-g). To study whether this rescue was detectable at the synaptic level, we performed electrophysiological analysis of the hippocampus and observed normalization of LTP in APP/PS1 animals upon augmented expression of ECSIT (Fig. 6i), with no significant change in basal transmission (Extended Data Fig. 7h). Taken together, these results show that increased ECSIT expression rescues cognitive deficits in the APP/PS1 mouse model of AD and demonstrate that maintenance of ECSIT expression is indeed protective against AD-like disease. This further corroborates the hypothesis that ECSIT downregulation observed in AD patient brains can contribute to disease progression.

## Discussion

In this study, we demonstrate that dysregulation of the mitochondrial protein ECSIT is associated with the development of AD and contributes to the development of AD-like pathology in animal models. An analysis of reported and predicted ECSIT interactions with redox and mitochondrial proteins encoded by AD susceptibility genes led to the hypothesis that ECSIT might be an interaction hub for many of these proteins, including PSEN1, and could thus play an important role in the etiology of AD^28, 29, 62, 63^. A role for mitochondrial dysfunction and mitochondrial ROS in AD had been suggested based on multiple studies, leading to the “mitochondrial cascade hypothesis”^4–7, 11, 33, 64^. However, the origin and nature of the mitochondrial dysfunction and aberrant mROS production has remained unclear ^16^. The role of ECSIT in CI stability^25, 27^, mROS production^26^ and mitophagy^27^ in immune cells led us to hypothesize that ECSIT protects against mitochondrial dysfunction and that neuronal ECSIT dysregulation might contribute to AD through increased mROS production, failed mitochondrial quality control and neuropathological oxidative stress leading to neuroinflammation and AD pathology.

In macrophages, ECSIT is critical for mitochondrial homeostasis in a broad range of metabolic contexts^27^. In contrast to macrophages, neurons are thought to be more reliant on OxPhos^3, 49, 65^ and should therefore exhibit more extensive metabolic disruptions and mROS production, and be especially sensitive to ECSIT dysregulation. Indeed, we observed significant mitochondrial dysfunction upon deletion of *Ecsit* in neurons. Deletion of *Ecsit* in only a subset of neurons led to the development of progressive neuropathology. This included abnormal cognitive behavior, neuronal loss, gliosis, neuroinflammation and tau pathology, mirroring the processes observed in human neurodegenerative diseases like AD. These results are consistent with recent reports of neurodegeneration in mouse models of mitochondrial dysfunction, such as *Mfn2* conditional KO^33^, *Drp1* conditional KO^66^ and *Aldh2* KO^67^ mice. However, none of these studies addressed whether mROS accumulation, as opposed to other consequences of CI disruption, directly triggered pathology.

In mice with reduced neuronal ECSIT expression, scavenging mROS with MCAT delayed neuropathology, implicating mROS derived from dysfunctional OxPhos as an etiopathological factor. Remarkably, other models of altered OxPhos in the CNS do not lead to age-related neuropathology: those either develop more rapidly, like the model of Leigh syndrome with *Ndufs4* KO^68, 69^; lack oxidative stress^70^; or have only mild effects on specific brain regions^71–75^.

Thus, the conditional *Ecsit* deletion in neurons of the cortex and hippocampus provides a unique model to address the role of CI-derived mitochondrial dysfunction and mROS in the pathogenesis of AD, in which CI dysfunction arises early^7^.

Mitochondrial dysfunction has been shown to appear early in the APP/PS1 murine model of AD^76^ and in AD patients^7^, and we found ECSIT to be downregulated in the hippocampus of young APP/PS1 mice. Further downregulation of ECSIT in the CNS of this AD mouse model resulted in worsening of the neuropathology: deficits in associative and spatial memory, synaptic dysfunction in hippocampus, decreased survival, and gliosis at the tissue level were all exacerbated. Consistent with a protective role for ECSIT, when we artificially elevated the expression of ECSIT in hippocampus of young APP/PS1 mice, these cognitive deficits were rescued.

Although neuronal mitochondrial dysfunction and increased mROS production should be addressed through mitochondrial quality control pathways, loss of neuronal ECSIT also resulted in a paradoxical disruption of mitophagy. ECSIT may therefore provide a link between the accumulation of mROS and defective mitophagy, which recent work has suggested to contribute to the development of AD pathology^21, 48^. We observed that ECSIT in neurons interacts with PINK1 and that lack of ECSIT prevents completion of mitophagy downstream of Parkin and LC3b recruitment to mitochondria. This observation, and our recent work in macrophages, suggests that ECSIT-dependent mitophagy may be an essential part of the basal mitochondrial quality control, as no acute mitochondrial damage was required to trigger this process^27^.

Surprisingly, impairing the classical pathway of damage-induced mitophagy in mouse models, via PINK1 and Parkin knock-out, failed to cause a widespread endogenous neuropathological phenotype^34, 77^, except when mitochondrial damage was induced at the same time^78, 79^. Another recent study suggested that basal mitophagy in neurons was independent of PINK1^49^. Nevertheless, changes in mitophagy observed upon ECSIT deletion, along with the alterations of PINK1 and Parkin expression we observed, are consistent with those seen in AD hippocampal tissue and neurons^22, 80^. Taken together, these results suggest a crucial role for ECSIT in a basal mitophagy pathway and maintenance of mitochondrial quality. As a result, decreased expression of ECSIT leads to mitochondrial dysfunction and mROS accumulation, *and* a failure of the neuronal mitochondrial quality control systems to correct this source of oxidative stress.

In summary, we propose that ECSIT dysregulation directly contributes to sporadic AD development and promotes worsening pathology. Regulation of ECSIT expression is poorly understood, and future work should examine transcriptional and post-translational regulation of ECSIT expression in the CNS. Although the *ECSIT* gene is located in a region linked to AD risk^28, 81^ and a polymorphism in the ECSIT interactant *NDUFS3* has recently been associated with a higher risk for AD^82^, so far no AD-associated polymorphism likely to affect ECSIT expression or function has been identified^80, 83, 84^. However, the location of *ECSIT* on chromosome 19 near *APOE* may have prevented identification of weaker risk factors^84^. A more precise study of the *ECSIT* genomic region might reveal intergenic polymorphisms associated with changes in its expression, or a combination of polymorphisms associated with significant risk^85, 86^. Additionally, interaction of ECSIT with enzymes involved in APP metabolism might be responsible for ECSIT downregulation and mitochondrial dysfunction. In the context of AD, PSEN1 could sequester or destabilize ECSIT outside of mitochondria, and contribute to its degradation, thereby affecting CI assembly. Indeed, PSEN1 mutations in familial AD have been suggested to function by trans-dominant inhibition of WT secretase complex^87^ and PSEN1 levels are elevated in sporadic AD^88^. An improved understanding of ECSIT regulation and the link between ECSIT-dependent mitophagy and neuropathological diseases will reveal new aspects of AD pathogenesis, and promises the identification of new therapeutic strategies for the treatment of AD in the future.

## Methods

### Mice

Sex- and age-matched mice were studied following experimental protocols approved by the Columbia University Institutional Animal Care and Use Committee and in accordance with the National Institutes of Health guidelines for animal care. Unless otherwise indicated, mice were group-housed in standard cages under a 12-h light/dark cycle with *ad libitum* access to water and food. The *Ecsit*-flox mouse strain has been described previously^27^. Surgical procedures were performed under anesthesia and all efforts were made to minimize suffering. Animals were housed individually following surgery. CamKII-Cre (JAX # 005359)^32^, Rosa26Cre-ERT2 (“ER-Cre”, from Dr. B. Reizis, made by Dr. T. Ludwig (unpublished and ^27^), and Synapsin-Cre (JAX # 003966) mouse strains were used to delete floxed *Ecsit* alleles in neurons at different ages. MCAT (JAX #016197) and APPswe/PS1dE9 (APP/PS1, MMRRC # 34832-JAX) mice were crossed to *Ecsit*-flox and Cre strains. It has come to our attention that breedings in which the sire carried CamKII-Cre or Synapsin-Cre transgene with a floxed *Ecsit* allele, could lead to the production of pups with germline recombination of the allele (represented by “-“). This has been reported by other investigators as well^89, 90^. Thus we avoided using Cre-carrying sires in breedings when possible and we designed a genotyping PCR to exclude from subsequent experiments animals in which germline recombination had occurred, when a Cre-carrying sire could not be avoided. In some figures, “CTRL” instead of “WT” represents a pool of CamKII-Cre^+^;ECSIT^+/+^ and CamKII-Cre^+^;ECSIT^+/-^ animals, after we verified that phenotypes were indistinguishable. C57BL/6 WT mice were purchased from The Jackson Laboratory.

### Primary cell preparation

Primary neuronal cell cultures were isolated from pooled hippocampi of neonatal (P0-P1) ER-Cre^+^;ECSIT^F/F^ pups or from the hippocampi of individual ER-Cre^+^;ECSIT^F/F^ or ER-Cre^+^;ECSIT^F/F^;MCAT^+^ pups. Brains were harvested in cold artificial cerebrospinal fluid (ACSF-1: 119 mM NaCl, 26.2 mM NaHCO_3_, 2.5 mM KCl, 1 mM NaH_2_PO_4_, 1.3 mM MgCl_2_, 10 mM D-glucose) and hippocampi dissected under a microscope in ACSF-1. Pooled hippocampi were digested in 0.25 % trypsin-EDTA (0.5 mL per pup) for 12 min at 37°C. DNAse-I (100 µg/mL) was added for 5 min at RT. Digestion was stopped with 3 washes by sedimentation with warm plating medium consisting of DMEM (Gibco) supplemented with 10 % heat-inactivated fetal bovine serum (FBS), penicillin (100 U/mL) and streptomycin (100 µg/mL). Tissues were dissociated in 1 mL of plating medium using a 1000 µL pipet tip, centrifuged at 200g for 5 min, resuspended in plating medium and filtered through a 0.45 µm cell strainer (Falcon). For individual pup neuronal preparation, a genotyping sample was kept and, after harvesting, tissues were processed similarly in a 1.5-mL tube using 0.5 mL 0.25 % trypsin-EDTA, DNAse-I and omitting the cell strainer step.

Cells were seeded into 6-well plates (0.4·10^6^ cells/well) or 24-well plates with glass coverslip (7.0·10^4^ cells/well), coated with 50 µg/mL poly-D-lysine (PDL, MP Biomedical cat# 102694) in 0.1 M boric acid, pH 8.5, for 16-24 h at 37°C. Cultures were maintained at 37°C in a humidified incubator with 5 % CO2. Six h later, medium was replaced with neurobasal medium supplemented with 2 % B27, 1 % glutamax, penicillin (100 U/mL) and streptomycin (100 µg/mL) (Gibco). Two µM cytosine arabinoside (AraC) was added after 2 days *in vitro* (DIV2). For inducible ECSIT deletion, 5 µM 4OH-tamoxifen dissolved in ethanol, or ethanol alone as control, was added at DIV2, 3 and 4. One half of the medium was replaced every 3 or 4 days. Neurons were analyzed at DIV7.

For neuron/astrocyte/microglia comparison by Western blot, single cell suspensions were made from cortices as described above and plated onto PDL-coated 6-well plates. The same culture protocol was followed for neuronal culture and mixed glia cultures were washed 3 times with PBS after 24 h and maintained in plating medium without AraC. At DIV11-14, mixed glia cultures were shaken at 100 rpm and 37°C for 2 h to detach microglia from astrocytes. Microglia were collected with a cold PBS wash, astrocytes were detached with 0.05 % trypsin-EDTA, and washed in plating medium. All cell cultures were washed twice in PBS before lysis. For astrocyte/microglia immunofluorescence, cells were detached at DIV14 and replated on PDL-coated coverslips in 24-well plates (1.0·10^4^ cells/well).

### Cell lines

293FT cells (Thermofisher) were plated in tissue culture dishes and maintained in standard culture medium composed of DMEM (Gibco) medium supplemented with 5 % FBS. Neuro2a cells were obtained from the ATCC® (CCL-131™) and maintained in Eagle’s Minimum Essential Medium (MEM) with 10 % FBS.

### Human participants

Human tissue obtained from the Columbia University Alzheimer’s Disease Research Center/New York Brain Bank was de-identified and as such is IRB-exempt under NIH IRB exemption four (E4), as described previously^91^. Frozen autopsy tissue from the cortex (frontal) of control subjects and subjects with severe AD were matched for age and post-mortem interval. Control subjects had no history of cognitive impairment and had no pathology that met the criteria for AD or any other age-related neuropathology. Subjects with severe AD had a history of cognitive impairment and Alzheimer’s pathology. Cortical samples were classified as “severe” based on a local assessment of the density of tau pathology^91^. Human brain tissues obtained from the Arizona Study of Aging and Neurodegenerative Disorders (AZSAND) database (www.brainandbodydonationprogram.org) were from the temporal cortex of individuals with Alzheimer’s disease and age-matched, non-Alzheimer’s disease controls. The diagnosis of AD using pathological criteria including Braak score is based on the description of Arizona Study of Aging and Neurodegenerative Disorders and Brain and Body Donation Program (Beach et al., 2015). Informed consent was obtained from these subjects. Detailed information for each of the cases studied is provided in table S1.

### Flow cytometry of primary neurons

At DIV7, neuronal culture medium was replaced with neurobasal medium without B27, which contains antioxidants, for 1 h. Rotenone (20 nM) or DMSO was added (Sigma-Aldrich) and incubated at 37°C for 15 min. Then 1 µM MitoSOX, a fluorochrome specific for anion superoxide produced in the inner mitochondrial compartment (Invitrogen), or the corresponding dilution of DMSO was added and further incubated at 37°C for 15 min. Mitotracker green FM (50 nM, Invitrogen) was added and incubated for 30 min at 37°C. Cells were washed with PBS, incubated in 0.5 % trypsin-EDTA for 2 min at 37°C, and collected in cold plating medium. Cells were centrifuged and resuspended in FACS buffer (PBS, 0.5 % FBS, 2.5 mM EDTA) with 1 µM DAPI, to allow for exclusion of dead cells (gating strategy in Extended Data Fig. 1f). Cells were then washed in FACS buffer and resuspended in FACS buffer. Cells were analyzed on a Fortessa or LSRII cytometer (BD) using FITC settings for mitotracker green and 585/42 emission filter with 488 nm laser. The corresponding mean fluorescence intensity of unstained controls was subtracted from the dye intensity and the fold over non-stimulated control was calculated.

### Mitophagy assay

Lentiviral shRNA constructs targeting GFP as a control (clone ID RHS4459) or ECSIT (clone ID TRCN0000113957) were described previously^26^ (Open Biosystems). The retroviral construct pCHAC-mt-mKeima (a generous gift from Richard Youle; Addgene plasmid # 72342) was used for mitochondrial expression of mKeima. This protein displays 2 fluorescence excitation/emission spectra distinguishable by flow cytometry and dependent on pH: “green” at neutral pH (ex 488/em 620 nm) and “red” (ex 521/em 610 nm) at acidic pH^92, 93^. Viral vector stocks were produced by calcium phosphate transfection of 293FT cells in combination with packaging vectors psPAX2 (a generous gift from Didier Trono; Addgene plasmid # 12260) for lentiviral constructs, or pCL-Eco (Addgene plasmid # 12371) for retroviral constructs, and envelope pCMV-VSV-G (Addgene plasmid #8454). Medium of 70 % confluent 293FT in 75-cm^2^ flasks was changed 2 h before transfection. Calcium phosphate precipitates were prepared by mixing 12.5 µg viral construct with 12.5 µg packaging plasmid and 5 μg pCMV-VSV-G in water for a final volume of 875 µL. Subsequently, 125 μL 2 M CaCl_2_ and 1 mL HBS 2X (50 mM HEPES, 10 mM KCl, 280 mM NaCl, 1.5 mM Na_2_HPO_4_, pH 7.05) were sequentially added dropwise in slowly vortexed solution. Solutions were incubated at RT for 20 min and added gently to 293FT supernatant. Medium was replaced by 7 mL of culture medium 24 h later. Supernatants were collected, centrifuged at 1,500 rpm for 5 min and filtered through a 0.45 µm filter (Pall). A total of 500,000 Neuro2a cells was incubated with 1 mL shRNA lentiviral vectors, 6 μg/mL polybrene (Sigma-Aldrich) and 10 mM HEPES (Invitrogen) for 3 h. Culture medium was added and replaced after 24 h. Fourty-eight h later, transduced Neuro2a cells were selected with 4 µg/mL puromycin for 72 h (Sigma-Aldrich) and maintained with 2 µg/mL puromycin. Neuro2a shGFP and shECSIT cells were then transduced with mt-mKeima retroviral vectors and populations of similar “green” fluorescence intensity were sorted by flow cytometry (FACSAria).

To assess the level of basal mitophagy, Neuro2a cells were plated on 12-well plates, treated or not with Bafilomycin A1 (10 nM, Sigma-Aldrich) for 16 h, to block lysosomal acidification, detached with 0.05 % trypsin EDTA and resuspended in cold FACS buffer. Green and Red fluorescence was analyzed by flow cytometry (BD Fortessa), using PE-TexasRed settings and 488 nm excitation with 610/20 emission filter, respectively. Bafilomycin condition was used as a reference to determine the percentage of mainly Red positive cells, undergoing active mitophagy. Green fluorescence was similar between cell types.

### Brain processing

Mice were sacrificed by cervical dislocation. Brain regions were dissected quickly in ACSF-1, snap frozen and stored at −80°C. Samples were homogenized by gentle, short sonication in 5 volumes of ice-cold homogenization buffer (10 mM tris-HCl pH 7.4, 140 mM NaCl) supplemented with 1 mM DTT and protease inhibitors (1 mM PMSF, 1 µg/mL aprotinin, 10 µg/mL pepstatin and 1 µg/mL leupeptin) and phosphatase inhibitors (2 mM sodium fluoride, 1 mM sodium orthovanadate, 1 mM β-2-glycerophosphate). Homogenates were aliquoted for protein, RNA and DNA preparation and processed or stored at −80°C. For histology, mice were euthanized with CO_2_ and transcardially perfused with 20 mL ice-cold PBS and 15 mL 4 % paraformaldehyde (PFA) in PBS at a rate of 5 mL/min. Brains were dissected and postfixed in 4 % PFA for 16 h. Coronal brain sections of 65 μm were produced using a vibrotome (Leica Microsystems).

### Mitochondria isolation

Mitochondria were prepared as described previously^94, 95^. Brains from 3- to 5-month-old mice were rinsed in ice-cold isolation buffer IBII (112.5 mM mannitol, 37.5 mM sucrose, 5 mM HEPES, 0.1 mM EDTA potassium pH 7.4) and cortices dissected, weighed, minced and IBII was added to obtain 100 mg tissue/mL. Tissue was homogenized with a Dounce homogenizer (Wheaton). A sample was saved as total tissue control, after clarification by centrifugation at 3000g for 30 min at 4°C. The resultant homogenate was diluted 1:2 in 24 % (vol/vol) Percoll prepared in IBI (112.5 mM mannitol, 37.5 mM sucrose, 5 mM HEPES, 1 mM EDTA potassium pH 7.4, 0.08 % fatty acid-free bovine serum albumin (BSA, Sigma-Aldrich)) and this 12 % Percoll solution layered on a discontinuous gradient containing 42 % and 24 % Percoll. This gradient was centrifuged at 27,000g for 10 min at 4°C. After centrifugation, band 2 (the interface between 12 % and 24 % layers containing synaptosomes) and band 3 (the interface between 24% and 42 % layers containing non-synaptosomal mitochondria) were removed from the density gradient. Both fractions were resuspended in 3 volumes of IBI and centrifuged at 16,700g for 10 min at 4°C. Pellets were resuspended in 300 µL IBII and centrifuged at 19,000g for 5 min at 4°C, washed once in ROS buffer (125 mM KCl, 2 mM K_2_HPO_4_, 5 mM MgCl_2_, 10 mM HEPES, 10 µM EGTA potassium, pH 7.4), then resuspended in ROS buffer and stored on ice. Protein concentrations were determined using the microBCA protein kit (Thermo Scientific). Ten μg synaptosomes were prepared in Laemmli buffer for Western blot analysis and 30 μg of synaptosomes were used for the mitochondrial potential assay.

For mitochondrial potential determination from permeabilized synaptosomes, 50 µg/mL digitonin was added to synaptosomes on ice, which were then plated in black 96-well plates in the presence of 5 mM complex I substrates glutamate and malate with or without 5 µM CCCP (carbonyl cyanide *m*-chlorophenyl hydrazone, Sigma-Aldrich) and 1 µM TMRM (tetramethylrhodamine, methyl ester, perchlorate; Invitrogen), and incubated at 37°C for 15 min. Fluorescence was read in a plate reader using 530 nm as excitation wavelength and 590 nm as emission wavelength (Biotek). Fluorescence from wells with buffers without mitochondria was subtracted and results are expressed as fold of non-treated WT.

### Amplex Red-based hydrogen peroxide measurements

Mitochondrial hydrogen peroxide production was measured by the Amplex Red hydrogen peroxide/peroxidase assay on non synaptosomal mitochondria in different substrate conditions, following manufacturer’s instructions (Molecular Probes). CII substrates (5 mM succinate) or CI substrates (5 mM glutamate and 5 mM malate) and antimycin (0.5 µM) or ROS buffer alone were plated in black 96-well plate. Amplex Red (50 µM) reagent containing 2 U/mL horseradish peroxidase was added next. Finally 5 µg mitochondria in ROS buffer or ROS buffer alone were added to the wells and incubated at 37°C. Fluorescence was kinetically followed during 20 min using 530 nm as excitation wavelength and 590 nm as emission wavelength (Biotek). Fluorescence from wells without mitochondria was subtracted and results are expressed as fold of WT.

### Determination of mitochondrial and intracellular ROS in brain

To estimate production of ROS, brain sections from 3- to 5-month-old mice were exposed to 2.5 µM MitoSOX Red (Molecular Probes), at 37°C for 30 min^14^. Images were captured by confocal microscopy as described^13, 96^. Quantification of staining intensity and the percentage of area occupied by MitoSOX was performed using Universal Image Program^13^. Evaluation of intracellular ROS levels was assessed by EPR (electron paramagnetic resonance) spectroscopy as previously described^14, 97^. Brain slices were incubated with CMH (cyclic hydroxylamine 1-hydroxy-3-methoxycarbonyl-2, 2, 5, 5-tetramethyl-pyrrolidine, 100 µM) and then washed with cold PBS. Tissues were harvested and homogenized with 100 µl of PBS for EPR measurement. The EPR spectra were collected, stored and analyzed with a Bruker EleXsys 540 x-band EPR spectrometer (Billerica, MA) using the Bruker Software Xepr (Billerica, MA).

### Coimmunoprecipitation (coIP)

50 µL of cortical homogenate from 3- to 5-month-old C57Bl/6 WT mice was diluted 20 times in coIP buffer (150 mM NaCl, 25 mM HEPES, 0.2 % NP40, 10 % Glycerol) with 1 mM DTT and protease inhibitors (1 mM PMSF, 1 µg/mL aprotinin, 10 µg/mL pepstatin and 1 µg/mL leupeptin), rotated for 10 min at 4°C and clarified at 3,000g for 30 min at 4°C. Lysate was incubated for 4 h at 4°C with anti-ECSIT2 antibody^39^ or IgG control (Cell Signaling, #2729), followed by overnight precipitation with 30 µL of washed Protein A sepharose 4B beads (Invitrogen). Unbound fraction was collected and beads were washed in coIP buffer and resuspended in 2X Laemmli buffer. Samples were fractionated by SDS-PAGE and analyzed by Western blotting.

### Immunofluorescent histology

Immunofluorescence stainings were performed on free-floating 65-µm sections. Sections were permeabilized in PBS containing 0.5 % triton X-100 for 30 min at room temperature (RT), then incubated in blocking buffer (PBS 0.5 % triton X-100 with 5 % normal goat serum and 5 % normal horse serum (Jackson Immunoresearch Laboratories)) for 2 h at RT. Sections were then incubated with primary antibodies in blocking buffer for 72 h at 4°C. The following primary antibodies were used: rabbit anti-ECSIT2 (dilution 1:100)^26^, mouse anti-GFAP (1:1,000; Sigma-Aldrich), rabbit anti-Iba1 (1:500; Wako), chicken anti-MAP2 (1:1,000; Abcam). Sections were washed 3 times with PBS 0.5 % triton X-100 for 30 min at RT and incubated with fluorophore-conjugated secondary antibodies (all 1:400) and NeuroTrace Deep-Red (1:250; Invitrogen) for 2 h at RT or 16 h at 4°C in PBS. After 3 washes in PBS for 30 min at RT, sections were mounted in Vectashield antifade mounting medium with or without DAPI (Vectorlabs) and covered with a #1.5 cover glass (Fisher Scientific). As negative control, rabbit IgG or mouse isotype control at the same concentration replaced primary antibody.

Confocal single plane tiled images of whole brain sections were taken with a Zeiss LSM 710 confocal microscope with four lasers (405, 488, 561, and 633 nm) and 10x, 40x or 63x objective lenses. Images were processed and quantified using Fiji (ImageJ). For quantification, sections from a comparable hippocampal position were used, with the Allen Adult Mouse Brain Atlas as a reference (http://mouse.brain-map.org/static/atlas,^98^). NeuroTrace-based analysis of structure thickness was measured at comparable locations perpendicularly to the axis of cortical or hippocampal layers, as represented in Extended Data Fig. 3j. NeuroTrace-based analysis of the area was measured by drawing the perimeter of brain regions according to the Allen Adult Mouse Brain Atlas. For MAP2 staining quantification, integrated intensity of the region of interest (delimited as described above) was divided by the mean intensity of an unaffected region of the same section comprising brain stem tissues. For GFAP- and Iba1-based quantification of gliosis, images were thresholded, converted to binary and the percentage of stained area was determined in each region of interest. Results are normalized to WT in each experiment. All quantifications were blinded.

### Immunofluorescence of primary neurons

Primary cells attached to coverslips were washed once with warm PBS, fixed in warm 4 % PFA at RT for 20 min, washed once in PBS and permeabilized with PBS containing 0.2 % Triton X-100 for 15 min. Cells were then blocked in PBS with 10 % FBS and 5 % normal goat serum for 1 h at RT. The following primary antibodies were prepared in blocking buffer (see above) and incubated for 2 h at RT: rabbit anti-ECSIT2 (1:100)^26^, chicken anti-MAP2 (1:3,000; Abcam), mouse anti-MTCOI (1:100; Abcam), mouse anti-TOM20 (1:200; Sigma-Aldrich), mouse anti-TOM6 (1:200; Santa Cruz Biotechnologies), mouse anti-TOM1 (1:100), rabbit anti-LC3b (1:200; Sigma-Aldrich), mouse anti-GFAP (1:4,000). Coverslips were washed 3 times with PBS and incubated with fluorophore-conjugated secondary antibodies diluted in PBS (all 1:200) for 1 h at RT. When indicated, Ib4-fluorescein (1:200; Vector Biolabs) or Acti-stain 488 phalloidin (100 nM, Cytoskeleton) was added to the secondary antibody solution. Coverslips were washed 3 times with PBS and mounted using Prolong gold with Dapi (Invitrogen). Cells were captured in at least 5 random fields using Zeiss LSM 710 confocal microscope and a 40x or 63x lens. As negative control, rabbit IgG or mouse isotype control at the same concentration replaced primary antibody. Neurons were identified by MAP2 positive staining and astrocytes by GFAP. Images were processed and quantified using Fiji ImageJ. Number of primary branches were determined using Sholl Analysis plugin of Fiji on MAP2 staining^31^. Mitochondrial length was measured in neurites on TOM20, TOM1 or MTCOI staining using Fiji. Approximately 5 cells per field and 3-6 fields per experiment were quantified. Percentage of mitochondria colocalizing with LC3b enrichment in neurites was determined on LC3b and MTCOI or TOM20 double staining.

Quantification was performed in a blinded fashion.

### Behavioral studies

Behavioral testing was conducted during the light phase of a 12-h light/dark cycle and consisted a battery of tasks carried out over a period of 2 weeks in the following order: 2-day radial arm water maze, contextual fear conditioning, cued fear conditioning, visible platform water maze, open field behavior and sensory threshold assessment, as described previously^99^. For MCAT rescue animals, the sequence of tasks was as follows: contextual fear conditioning, cued fear conditioning, open field behavior and sensory threshold assessment.

### Open field behavior

Animals were placed into a plexiglass chamber (27.3 cm long × 27.3 cm wide × 20.3 cm high) for 10 min on each of two successive days during which time their movements were tracked using arrays of infrared beams and a computerized tracking system. Their movements were then analyzed using behavioral analysis software (Med Associates).

### Radial arm water maze (RAWM)

Testing was performed in a 120-cm-diameter pool containing a six-arm radial maze insert and filled with opaque water as described previously^100^. Mice were tested in fifteen 1-min trials on each of 2 consecutive days. The location of the escape platform was held constant during testing but the start location was pseudorandomly varied throughout. On the first day, training alternated between visible and hidden platform trials, while on the second day only hidden platform trials were conducted. Water temperature was maintained at approximately 24°C and mice were dried and placed in a clean heated cage between trials to prevent hypothermia. Entries into maze arms that did not contain the escape platform were scored as errors. Data are presented as the average number of errors committed during blocks of 3 training trials.

### Visible platform water maze

This task was conducted in the same 120-cm-diameter pool used for the RAWM task but with the partitions removed. Training for this task was carried out over 2 days with 3 morning and 3 afternoon trials on each day. Intertrial intervals were 15 to 20 min and rest periods between morning and afternoon sessions were 2–3 h. Each trial was a maximum of 60 s during which time the animals were required to swim to a visible escape platform located just above the water surface. Animals that did not reach the platform within the allotted time were guided to it and allowed to sit there for 15 s before returning to their home cage. The location of the platform was varied among 4 different locations such that it was not present in the same location on any two successive trials. Water temperature was maintained at approximately 24°C, and animals were dried and placed in a clean warmed cage after each trial to prevent hypothermia. Animal movements were recorded using a video-tracking system and time required to reach the platform (latency) and swim speed were determined using Ethovision XT behavioral analysis software (Noldus).

### Fear conditioning

Animals were placed into a conditioning chamber (30 cm x 30 cm x 30 cm) made of transparent Plexiglas (Noldus). The top unit of the chamber had a yellow light together with a video camera connected to a personal computer. Foot shocks were administered through a removable 32-bar grid floor and the entire apparatus was cleaned and deodorized between animals with distilled water and 70% ethanol. Animals were placed in the conditioning chamber once on each of three consecutive days. On the first day of exposure mice were placed in the conditioning chamber for 2 min before the onset of a discrete 30 s, 2,800 Hz, 85 dB tone (conditioned stimulus (CS)), the last 2 s of which coincided with a 0.8 mA foot shock (unconditioned stimulus (US)). After the CS/US pairing, the mice were left in the conditioning chamber for another 30 s before returning to their home cages. Twenty-four h after their first exposure, animals were returned to the conditioning chamber for 5 min without foot shock or tone presentation for contextual fear learning evaluation. During the third day, we evaluated the cued fear learning. Mice were placed in a novel context (rectangular black cage with vanilla odorant) for 2 min (pre-CS test), after which they were exposed to the CS for 3 min (CS test), and freezing was measured. Freezing behavior during all phases of testing was calculated using Ethovision XT software (Noldus).

### Sensory threshold assessment

Animals were placed into the same apparatus used for contextual fear conditioning. A sequence of single, 1-s foot shocks were then administered at 30-s intervals and 0.1-mA increments from 0 mA up to a maximum of 0.7 mA. Each animal’s behavior was monitored by the experimenter to determine their thresholds for first visible response to the shock (flinch), their first gross motor response (run/jump), and their first vocalized response.

### Hippocampal electrophysiological studies

Recordings of extracellular field potential were performed on acute hippocampal slices prepared as described previously^101^. Brains were rapidly removed and cooled in ice-cold ACSF-2 consisting of 124 mM NaCl, 4.4 mM KCl, 1 mM Na_2_HPO_4_, 25 mM NaHCO_3_, 2 mM CaCl_2_, 2 mM MgCl_2_, and 10 mM glucose. Hippocampi were then dissected and sliced into 400 µM sections using a tissue chopper. Slices were incubated at 29°C in an interface chamber under continuous perfusion (2 mL/min) with ACSF-2 continuously bubbled with 95% O_2_ and 5% CO_2_ and allowed to recover for a minimum of 90 min prior to recording responses in the CA1 region to stimulation of Schaffer collateral projections with a bipolar electrode. After evaluation of basal synaptic transmission, input/output relationships were determined prior to each recording and stimulus intensities that elicited 30% of the maximal response were utilized. Stable baselines were obtained for a minimum of 15 min and a theta-burst stimulation protocol consisting of 3 trains separated by 15-s intervals with each train consisting of 10 bursts at 5 Hz and each burst consisting of 5 pulses at 100 Hz was used to elicit LTP. PPF was elicited through a double stimulation delivered at different interstimulus intervals (ranging from 10 to 1,000 ms) and then comparing the ratio between the slope of the evoked response after the second stimulus and the slope of the evoked response from the first stimulus.

### Basolateral amygdala (BLA) electrophysiological studies

Mice were transcardially perfused with carbogenated ice-cold ACSF-3 (118 mM NaCl, 10 mM glucose, 2.5 mM KCl, 1 mM NaH_2_PO_4_, 1 mM CaCl_2_, and 1.5 mM MgSO_4_ (325 mOsm, pH 7.4)) for 60 s under pentobarbital anesthesia. Brains were removed, and 300-µm coronal slices containing the amygdala were prepared on a vibratome. Slices were allowed to recover for at least 1 h before being transferred to a recording chamber maintained at 32°C and perfused with the same ACSF-3 supplemented with 2 mM CaCl_2_. BLA pyramidal neurons were visualized using a 40x water-immersion objective and video-enhanced differential interference contrast microscopy on an upright microscope. Spontaneous excitatory postsynaptic currents (sEPSCs) were recorded at −60 mV in the presence of bicuculline. Excitatory post-synaptic currents (EPSCs) were recorded in the presence of bicuculline by stimulating the BLA using an extracellular electrode. The AMPA/NMDA ratio was calculated by dividing the peak EPSC at −60 mV by the NMDA receptor current measured at 50 ms after the peak at 40 mV. The composition of the electrode solution was 130 mM CsCl, 10 mM HEPES, 0.5 mM EGTA, 5 mM QX-314, 0.2 mM GTP, and 5 mM MgATP (osmolarity 290-300 mOsm, pH 7.4). Series resistance was monitored throughout the experiment. fEPSPs were recorded from BLA using a glass electrode filled with 2 M NaCl and by local stimulation with a concentric bipolar electrode.

Bicuculline was included in the recording solution. LTP was induced by a tetanic stimulation, 100 Hz for 1 s, repeated 4 times with an interval of 20 s^102^. Data were acquired using a Multiclamp 700B amplifier connected to a Digidata 1550A (Molecular Devices). Clampfit (Molecular Devices) and Mini Analysis programs were used for data analysis.

### Processing of human brain samples

Frozen fragments of human cortex were homogenized in RIPA buffer (50 mM Tris-HCl pH 7.4, 150 mM NaCl, 0.5 % sodium deoxycholate, 1 % NP40) with 1 mM DTT, protease and phosphatase inhibitors, and rotated for 30 min at 4°C. Samples were centrifuged for 30 min at 15,000 rpm for 30 min before collection of supernatants. For mRNA quantification, frozen fragments of human cortex were homogenized with a motorized hand pestle in RLT lysis buffer (Qiagen) and qPCR was performed as described in qPCR section.

### Western blotting

Cells in culture were washed twice in PBS and collected in triton lysis buffer (1 % Triton X-100, 150 mM NaCl, 50 mM HEPES pH 7.5, 5 mM EDTA) with 1 mM DTT and protease inhibitors (1 mM PMSF, 1 µg/mL aprotinin, 10 µg/mL pepstatin and 1 µg/mL leupeptin) and incubated for 20 min on ice. Lysates were then clarified at 15,000 rpm for 15 min at 4°C. For protein lysate preparation from brain homogenates, an equal volume of modified 2X RIPA buffer (80 mM tris-HCl pH 7.4, 2 % NP-40, 2 mM EDTA, 0.5 % sodium deoxycholate) with 1 mM DTT and protease and phosphatase inhibitors was added to homogenate and incubated on ice for 20 min. Lysates were centrifuged at 3,000g for 30 min at 4°C and supernatants collected. Protein concentrations were determined using the microBCA protein kit (Thermo Scientific). Ten to 20 µg total cell or brain lysate, mitochondrial or synaptosomal fractions were prepared in Laemmli buffer and separated by SDS-PAGE before transfer to a PVDF membrane (Immobilon-P, Millipore). Membranes were blocked in 5 % non-fat milk in TBS-Tween 20 (0.1 %) and incubated with primary and horseradish peroxidase (HRP)–conjugated secondary antibodies. We used Super Signal West Pico chemiluminescent substrate (Thermo Fisher Scientific) or Luminata Forte (Millipore) and BioExcell autoradiographic films (WWMP) for image development. Molecular weight from standard ladder is indicated in kDa and represented by a dash. When no ladder band was in proximity to the protein band, apparent band size is indicated, without dash. Band intensities were measured using Fiji ImageJ and normalized to a loading control, as indicated in figure legends.

Primary antibodies were: anti-VDAC1 (dilution 1:2,000, Abcam, clone 20B12AF2), NDUFS3 (1:1,000, Abcam, clone 3F9DD2), SDHA (1:2,000, Abcam, clone EPR9043), UQCRC2 (1:2,000, Abcam, clone 13G12AF12BB11), MTCOI (1:4,000, Abcam, clone 1D6E1A8), NDUFAF1 (1:2,000, Origene), TOM20 (1:500, Sigma-Aldrich, clone 4F3), HSP60 (1:500, Santa Cruz Biotechnology, clone H-1), GAPDH (1:5,000, Fitzgerald), beta-Tubulin (1:1,000, Sigma-Aldrich, clone TUB2.1), Actin (1:1,000, Santa Cruz Biotechnology, clone C-4), GFP (1:1,000, Santa Cruz Biotechnology, clone B-2), Parkin (1:1,000, Cell Signaling, Prk8), PINK1 (1:1,000, Novus Biologicals), LC3b (1:2,000, Sigma-Aldrich), 4-HNE (1:500, R&D, clone 198960), GFAP (1:1,000, Sigma-Aldrich, clone G-A-5), MAP2 (1:1,000, Millipore, clone AP20), PSEN1 (1:500, Santa Cruz Biotechnology, clone H-5), tau (1:500, Invitrogen, clone TAU-5), tau (1:2,000, Dako), TOM20 (1:1,000, Sigma-Aldrich), anti-phosphorylated tau CP13 (phosphoserine 202) and PHF1 (phosphoserine 396/404) (both 1:500 and a generous gift from Dr. P. Davies), human ECSIT (1:1,000, Lifespan), and rabbit polyclonal antibody against mouse ECSIT2 was previously described (1:1,000)^24, 26^.

### Quantitative PCR (qPCR)

For RNA isolation from mouse brain samples, 100-50 μL of homogenate was added to 600 μL of RLT buffer with β-mercaptoethanol and RNA isolated using QIAshredder and the RNeasy Mini Kit following the manufacturer’s instructions for tissue samples (Qiagen). RNA purity and quantity were analyzed by photometry (Gen5 BioTek).

For cDNA synthesis, 1-2 µg total RNA were reverse transcribed into cDNA using the Superscript III enzyme and oligo(dT) primers (Invitrogen). qPCR was performed using SYBR Green (Quanta). Primer sequences are listed in the tables below. Differences in cDNA inputs were corrected by normalization to HPRT cDNA levels for mouse and RPLP0 cDNA for human samples. Relative quantitation of target cDNA was determined by the formula 2^-ΔCT^, with ΔCT denoting fold increases above respective controls.

### mtDNA Mitomass

For DNA isolation from brain samples, PBS was added to 50-15 µL of homogenate for a total volume of 200 µL and the DNeasy Blood and Tissue kit was used, following manufacturer’s instructions for cell pellets (Qiagen). Genomic DNA concentrations were determined by photometry (Gen5 BioTek). Fifteen ng, 7.5 ng and 3.75 ng DNA were used to perform qPCR targeting the mitochondrial gene *mtCOI* and the nuclear gene *Ndufv1*. Ratios of 2^-CT^ for *mtCOI* over *Ndufv1* for the different DNA concentrations were averaged and fold of WT is shown.

### List of mouse primers used for qPCR

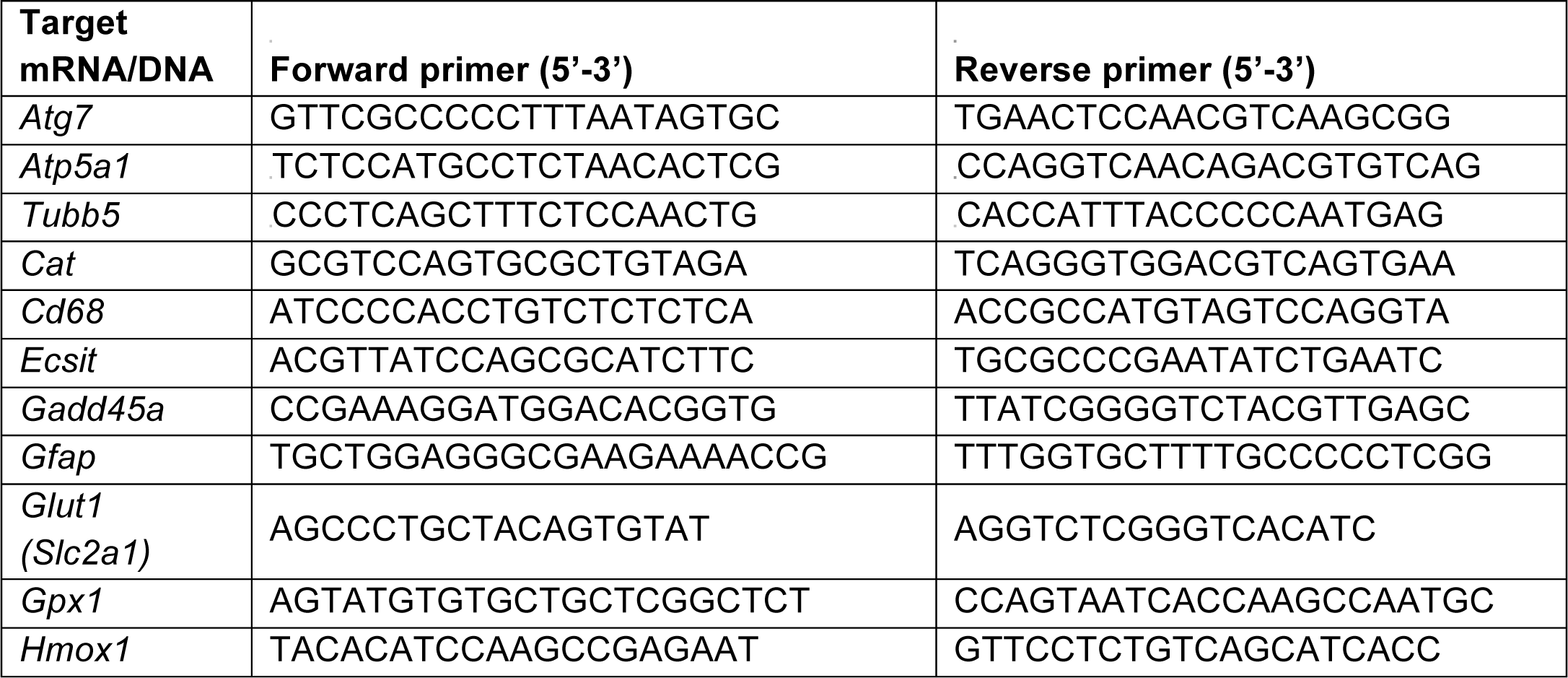
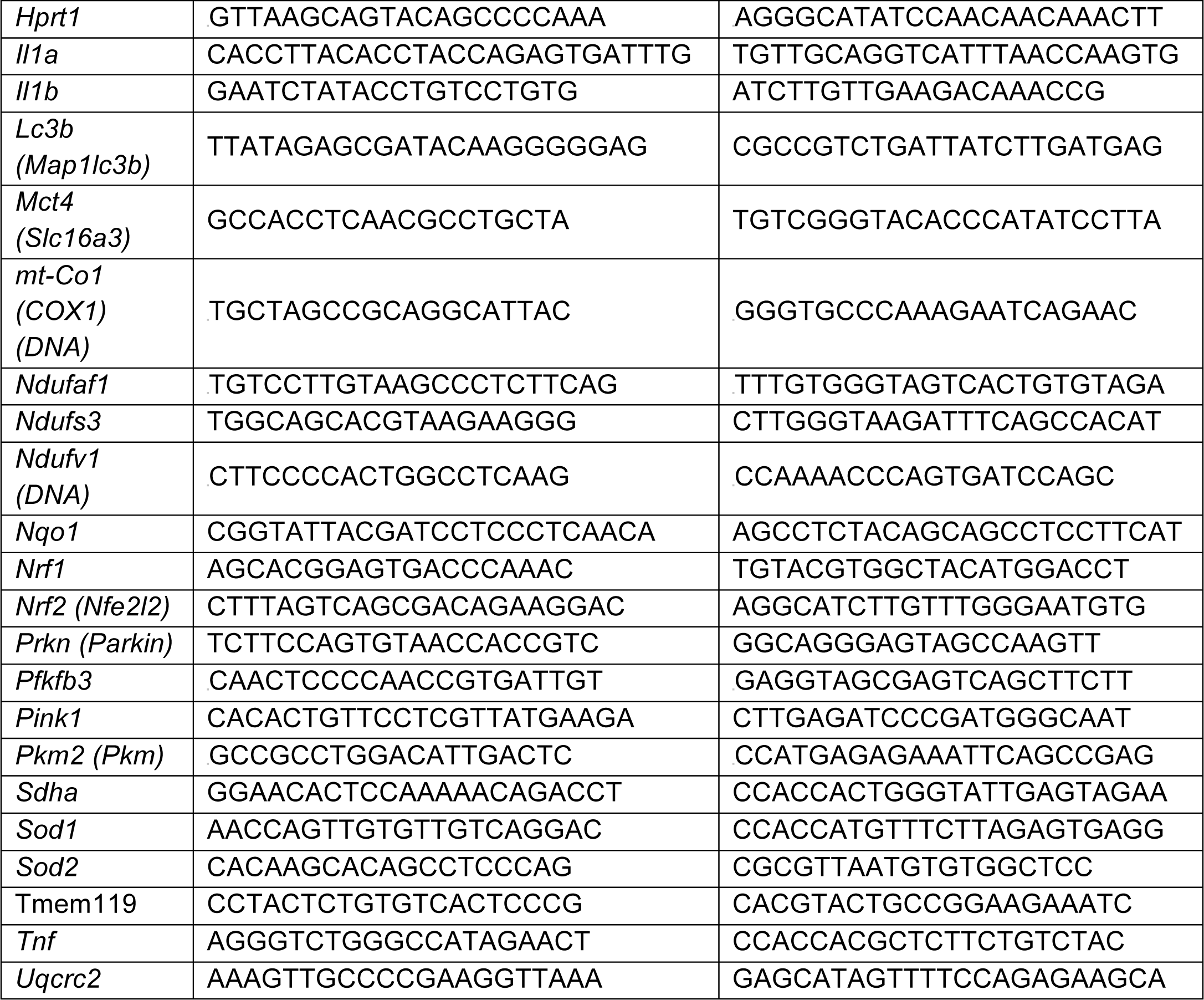

### List of human primers used for qPCR

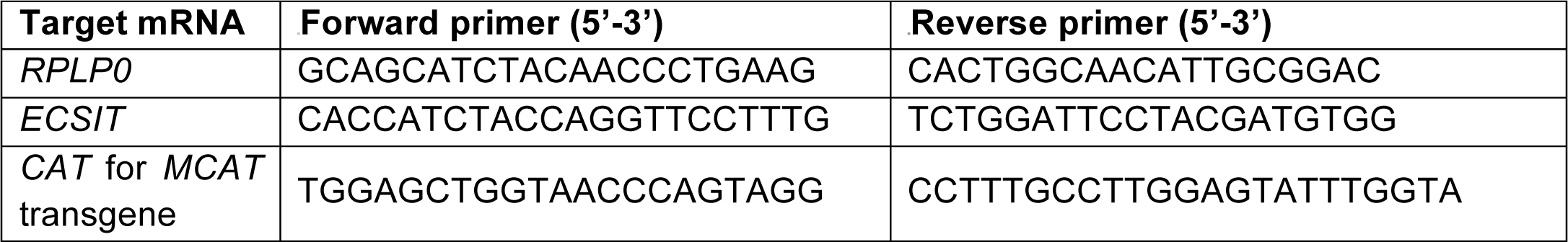

### Aβ ELISA

Brain homogenates were incubated in 5 M guanidine-HCl and 50 mM Tris-HCl pH 8.0 overnight at 4°C, diluted with 9 volumes PBS with protease inhibitors and clarified for 20 min at 16,000g and 4°C. Samples were further diluted 1:100 in standard diluent and Aβ levels were measured with the human Aβ1–42 ELISA kit (Invitrogen) following the manufacturer’s instructions.

### AAV injection

Male and female 2-month-old APPswe/PS1dE9 mice and littermate controls were used for intrahippocampal stereotaxic injections. Mice were deeply anesthetized with 500 mg/kg Avertin (2,2,2-tribromoethanol) and placed in the stereotaxic frame. AAV-GFP (AAV9-hSYN1-eGFP-WPRE, Vector Biolabs) and custom-made AAV-ECSIT-GFP (AAV9-hSYN1-eGFP-2A-mECSIT-WPRE, with eGFP and mouse ECSIT2 expression driven by the same hSYN1 promoter, with a T2A peptide linking eGFP and ECSIT, Vector Biolabs) were injected bilaterally into the dorsal hippocampus at the following coordinates (flat skull position): antero-posterior: −1.9 mm, medio-lateral: +/-1.8mm, dorso-ventral: −1.8 mm below dural surface as calculated relative to bregma according to the stereotaxic atlas. For each AAV, 0.5 µL/side (≈ 2.35×10^10^ genome copies) were injected with a 5-µL Hamilton syringe at a rate of 0.25 µL/min by a nano-injector system. The needle was allowed to remain in the brain for an additional 1 min before it was slowly retracted. Three months post-injection, mice were subjected to behavioral testing and electrophysiological analysis.

### AAV infection of primary neurons

Primary neurons were infected with 1.5×10^6^ GC/cell of AAV-GFP and AAV-ECSIT-GFP at DIV3 and fixed for immunofluorescence staining at DIV 10.

### Analysis of publicly available data

We analyzed the expression of ECSIT in GSE36980 from the Hisayama study. Human brain samples classified into healthy (n=47) and AD (n=32) were processed by array (Affymetrix Human Gene 1.0 ST Array)^53^. We compared log2 fold changes in pooled brain regions (hippocampus, temporal cortex, frontal cortex). We checked the expression of ECSIT and other indicated genes in GSE5281: gene expression in microdissected healthy neurons from entorhinal cortex from AD Braak V-VI patients (n=18) and controls (n=15) was measured by array (Affymetrix U133 Plus 2.0 array)^54^. We retrieved log2 fold change (LogFC) and adjusted p-values.

### Statistics

Statistical analysis was conducted using GraphPad Prism 6 software. All data were tested for normal distribution of variables. All normally distributed data were displayed as means ± standard error of the mean (sem) unless otherwise noted. Comparisons between two groups were performed with an unpaired Student’s t-test if normally distributed, or Mann-Whitney U test if otherwise. Groups of three or more were analyzed by analysis of variance (ANOVA) or the Kruskal-Wallis test, with Sidak’s or Dunn’s post-hoc test respectively. All statistical tests were two-tailed. Sample sizes for each experiment are reported in the figures and figure legends. A p-value of less than 0.05 was considered significant. Statistical parameters for each experiment can be found within the corresponding figure legends.

## Acknowledgements

**Funding:** This work was supported by NIH grant R56AG058449 to SG and OA. S.S.Yan were supported by NIH grant R37AG037319. Alice Lepelley was supported by a fellowship from BrightFocus foundation (A2014425F). We are grateful to Cristian Boboila and Erica Acquarone for technical assistance and Chung Dang for critical reading of the manuscript. We thank Qing Yu and Shijun Yan for assistance in mitosox experiment. We thank Dr. Peter Davies (Albert Einstein School of Medicine, NY) for the generous gift of anti-tau antibodies. Plasmid pCHAC-mt-mKeima was a gift from Richard Youle (Addgene plasmid # 72342), pCL-Eco a gift from Inder Verma (Addgene plasmid # 12371). We are grateful to the New York Brain Bank and Columbia University Alzheimer’s Disease Research Center (funded by NIH grant P50AG008702 to S.A. Small (P.I.) and Jean-Paul Vonsattel, Etty Cortes and L.S. Honig for the generous gift of post-mortem samples and associated information. We thank Thomas G. Beach and Geidy E. Serrano of the Banner Sun Health Research Institute Brain and Body Donation Program (BBDP, supported by National Institute of Neurological Disorders and Stroke, U24 NS072026 National Brain and Tissue Resource for Parkinson’s Disease and Related Disorders; National Institute on Aging, P30 AG19610 Arizona Alzheimer’s Disease Core Center; Arizona Department of Health Services, contract 211002, Arizona Alzheimer’s Research Center; Arizona Biomedical Research Commission, contracts 4001, 0011, 05-901 and 1001 to the Arizona Parkinson’s Disease Consortium; Michael J. Fox Foundation for Parkinson’s Research) for providing human brain tissues.

## Author contributions

Conceptualization, A.L., O.A., and S.G.; Methodology, A.L., A.F.T., I.N., S.S.Y., O.A. and S.G.; Investigation, A.L., L.V., A.S., H.Z., F.D., P.K., and Z.T.; Resources, F.R.G.C., A.F.T., Writing – Original Draft, A.L., O.A. and S.G.; Writing – Review and Editing, A.L., O.A, M.S.H., T.S.P. and S.G.; Supervision, I.N., S.S.Y., O.A. and S.G. Funding Acquisition, A.L., O.A. and S.G.

## Author information

The authors declare no competing interest.

---

**Extended Data Fig. 1.**
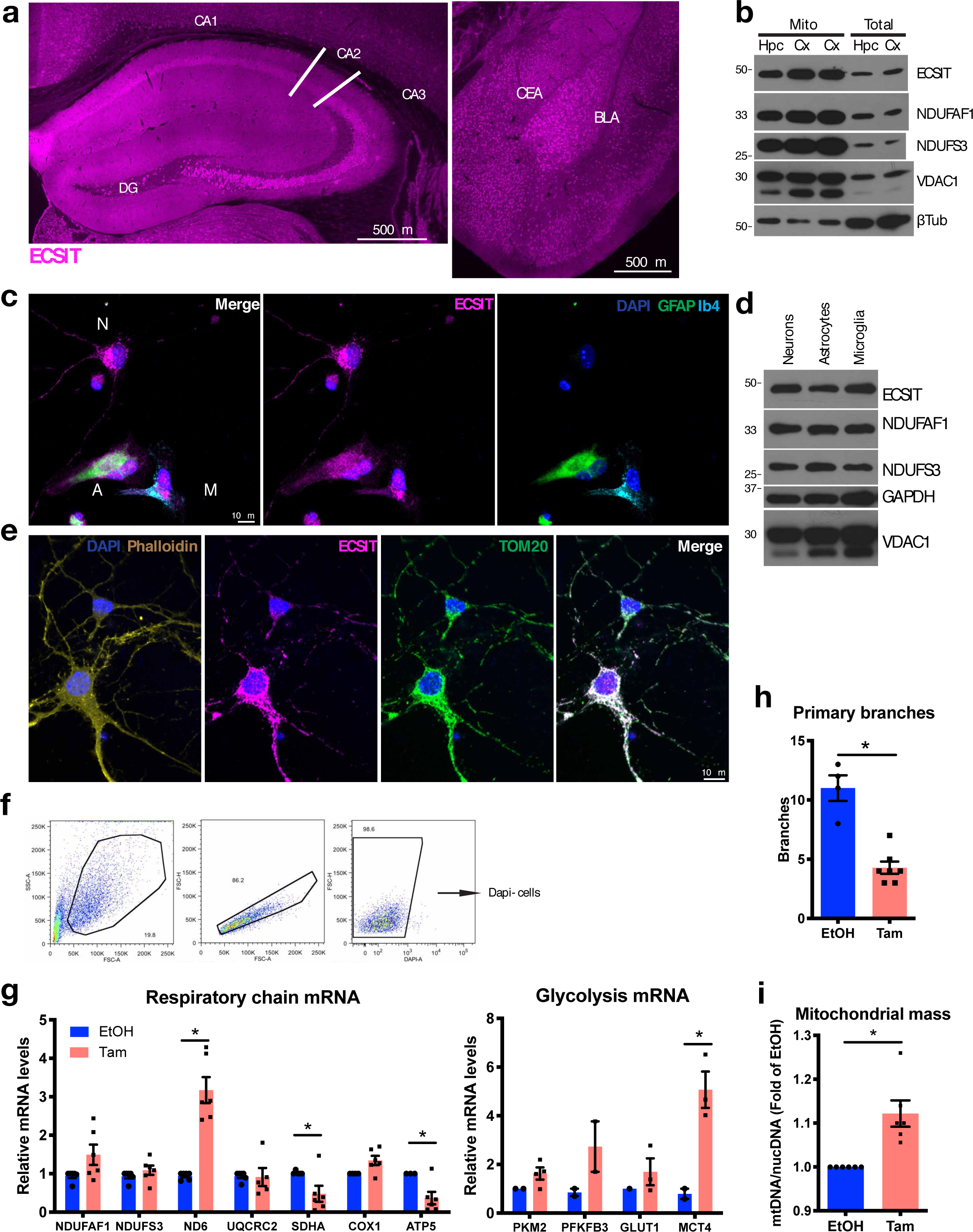
Loss of ECSIT in neurons leads to oxidative stress. **a,** ECSIT staining of coronal section from a 3-month-old WT mouse and confocal acquisition. (Left) Enlargement of hippocampus: areas CA1, CA2, CA3 and dentate gyrus (DG) are indicated. (Right) enlargement of amygdala region: central amygdala (CEA) and basolateral amygdala (BLA) are indicated. **b,** Levels of ECSIT and indicated mitochondrial proteins in mitochondria isolated from cortex (Cx) and hippocampus (Hpc) assessed by Western blotting of lysates from a 3-month-old WT mouse, representative of 4 independent experiments. **c,** Immunofluorescence staining and confocal acquisition of ECSIT in DIV10 primary mixed cultures of astrocytes (A, GFAP+), microglia (M, Ib4+) and neurons (N). **d,** Levels of ECSIT, NDUFAF1, complex I NDUFS3 and mitochondrial porin VDAC1 in lysates from astrocytes and microglia from primary mixed glia cultures and DIV10 primary neurons by Western blot. Representative of 3 independent experiments. **e,** Immunofluorescence staining and confocal acquisition of ECSIT, mitochondria (TOM20) and actin (phalloidin) in DIV10 primary neurons. Representative of 5 independent experiments. **f,** Gating strategy for neurons, Dapi-cells were considered in Fig 1d-e. **g,** mRNA levels of respiratory chain subunit (n=6) and glycolysis (n=3) genes in DIV7 primary neurons by qPCR. Average of experiments is shown; *, p<0.05 in multiple t-test. **h,** Sholl analysis with Fiji software of DIV7 primary neurons. Ten cells of each group were analyzed blindly; *, p<0.05 in t-test. Representative of 5 independent experiments. **i,** Mitochondrial mass assessed by the ratio of mitochondrial genomes (*mtCOI* gene) copy numbers over nuclear genomes (*Ndufv1* gene) by qPCR in primary neurons. Data are expressed as fold-increase relative to EtOH. Average of n=6 experiments; *, p<0.05 in Mann-Whitney test.

**Extended Data Fig. 2.**
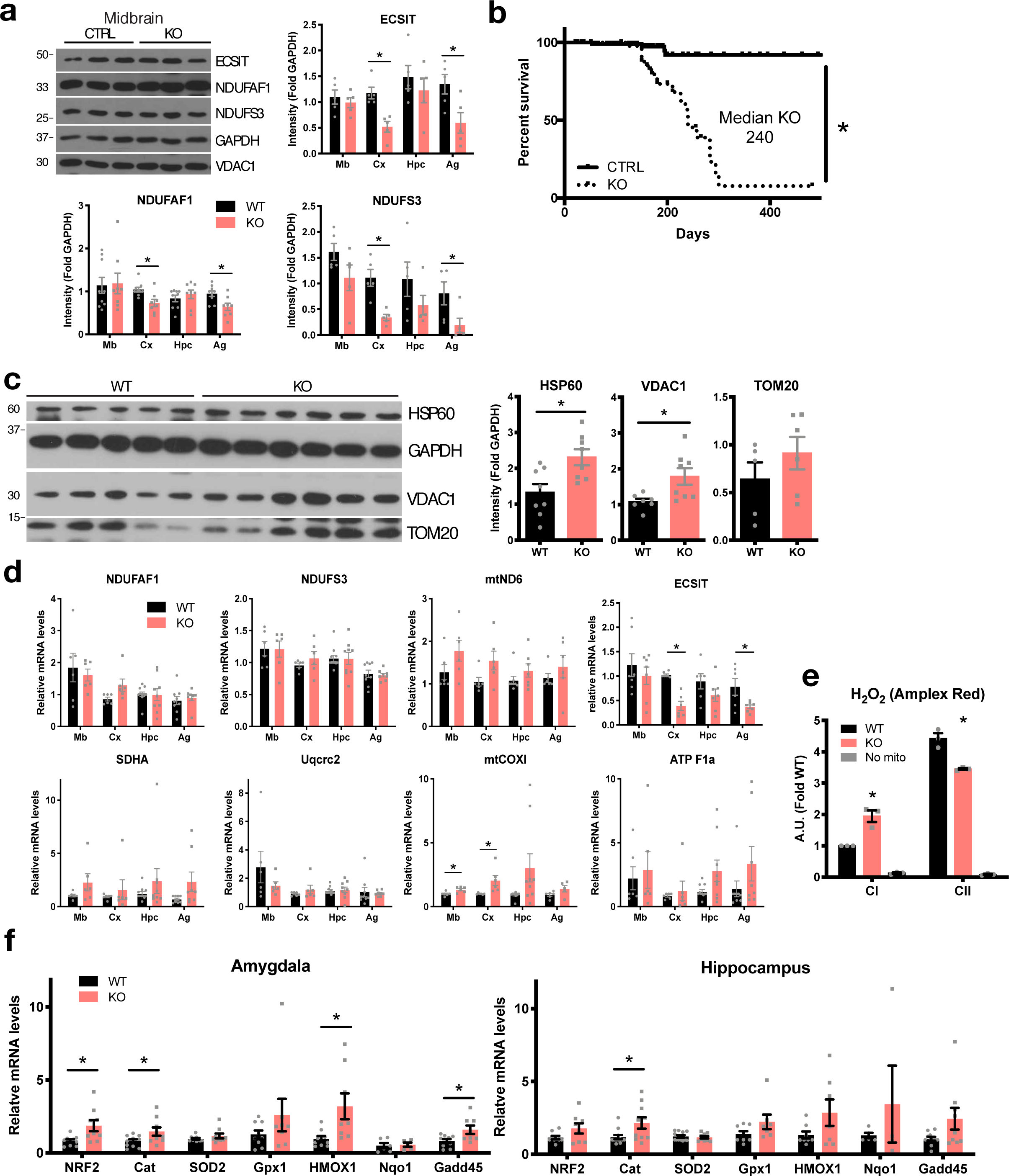
Deletion of ECSIT in neurons *in vivo* leads to mitochondrial dysfunction and oxidative stress in the brain. **a,** (Top left) Western blot analysis of the levels of ECSIT, Complex I (NDUFS3, NDUFAF1) and mitochondrial porin (VDAC1) in midbrain lysates. Each lane represents an animal. Representative of 3 independent experiments. (Right and below) Quantification of Western blot analysis for ECSIT, complex I (NDUFS3, NDUFAF1) and mitochondrial porin (VDAC1) in brain lysates. N=5-8 animals per group, *, p<0.05 in multiple t-test. **b,** Survival of controls (CTRL, n=100) and CamKII KO animals (n=47), *, p<0.05 in Mantel-Cox test. **c,** Western blot analysis of mitochondrial proteins HSP60, TOM20 and VDAC1 in cortex lysates. Quantification of band intensities over GAPDH on the right. N=5-8 animals per group, *, p<0.05 in t-test. **d,** mRNA levels of respiratory chain subunit genes in brain lysates by qPCR. Genes tested encode subunits of complex I (NDUFAF1, NDUFS3 and mtND6), complex II (SDHA), complex III (Uqcrc2), complex IV (mtCOXI) and complex V (ATP F1a). N=6 animals per group. *, p<0.05 in multiple t-test. **e,** H_2_O_2_ levels measured by Amplex Red fluorescence in cortical mitochondria isolated from animals of indicated genotypes and incubated in CI or CII substrates. n=3 experiments with 1 animals per group each. *, p<0.05 in 2-way ANOVA with Sidak’s post-hoc. **f,** mRNA levels of antioxidant response genes in hippocampus and amygdala by qPCR. N=9 animals per group. Data are expressed as average of fold increase relative to WT in each brain region. *, p<0.05 in multiple t-test. All experiments were performed with 3- to 5-month-old mice.

**Extended Data Fig. 3.**
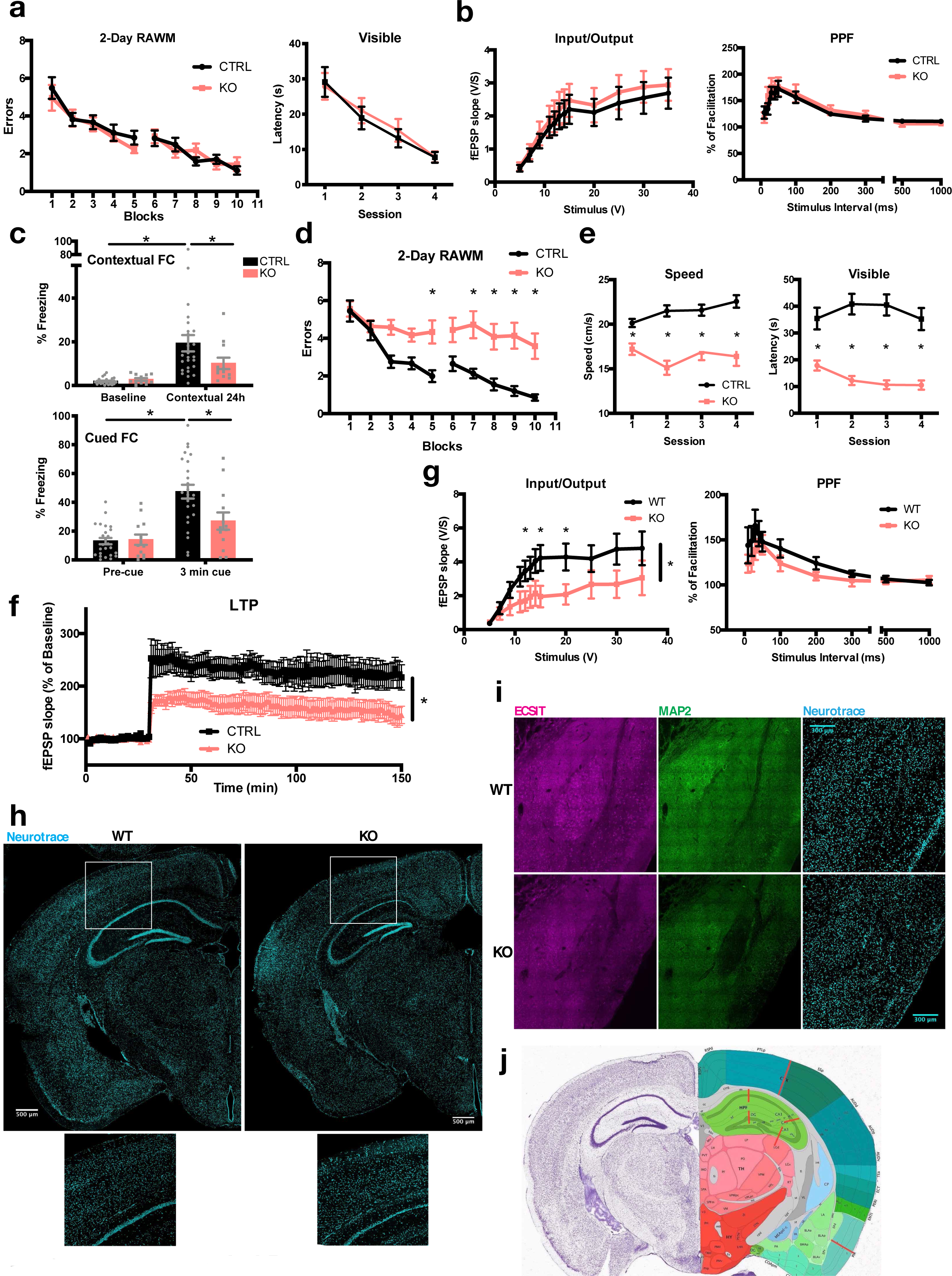
ECSIT deletion in neurons leads to cognitive dysfunction and neurodegeneration. **a,** (Left) Number of errors per block of trials in 2-day Radial Arm Water Maze (RAWM) in 3- to 5-month-old animals. (Right) visual control; n=9 CTRL; 9 KO. **b,** Basal synaptic transmission (Input/Output) and paired-pulse facilitation (PPF) in hippocampus of 3- to 5-month-old animals; n=11 CTRL; 6 KO. **c,** Fear conditioning (FC) in 6- to 8-month-old animals; n=25 CTRL; 12 KO. *, p<0.05 in Mann-Whitney test. **d,** Two-day RAWM with animals shown in **c**, *, p<0.05 in 2-way ANOVA with repeated measures with Bonferroni post hoc test. **e,** Visual, motor and motivational controls for animals shown in **c**. *, p<0.05 in 2-way ANOVA with repeated measures and Sidak post hoc test. **f,** LTP in hippocampus in 7- to 10-month-old animals (n=12 CTRL; 11 KO). *, p<0.05 in ANOVA with repeated measures after tetanus. **g,** Basal synaptic transmission and PPF in slices shown in **f**. *, p<0.05 in ANOVA with repeated measures. **h,** Representative Neurotrace staining of coronal sections of 8-month-old animals. **i,** Representative immunofluorescence staining of MAP2 and confocal acquisition in amygdala of 8-month-old animals. **j,** Reference coronal section taken from the Allen Brain Atlas used to measure thickness in Fig. 3f. Red bars represent location of thickness measures. For hippocampus, the thickness of the pyramidal layer cut by the red bar was considered.

**Extended Data Fig. 4.**
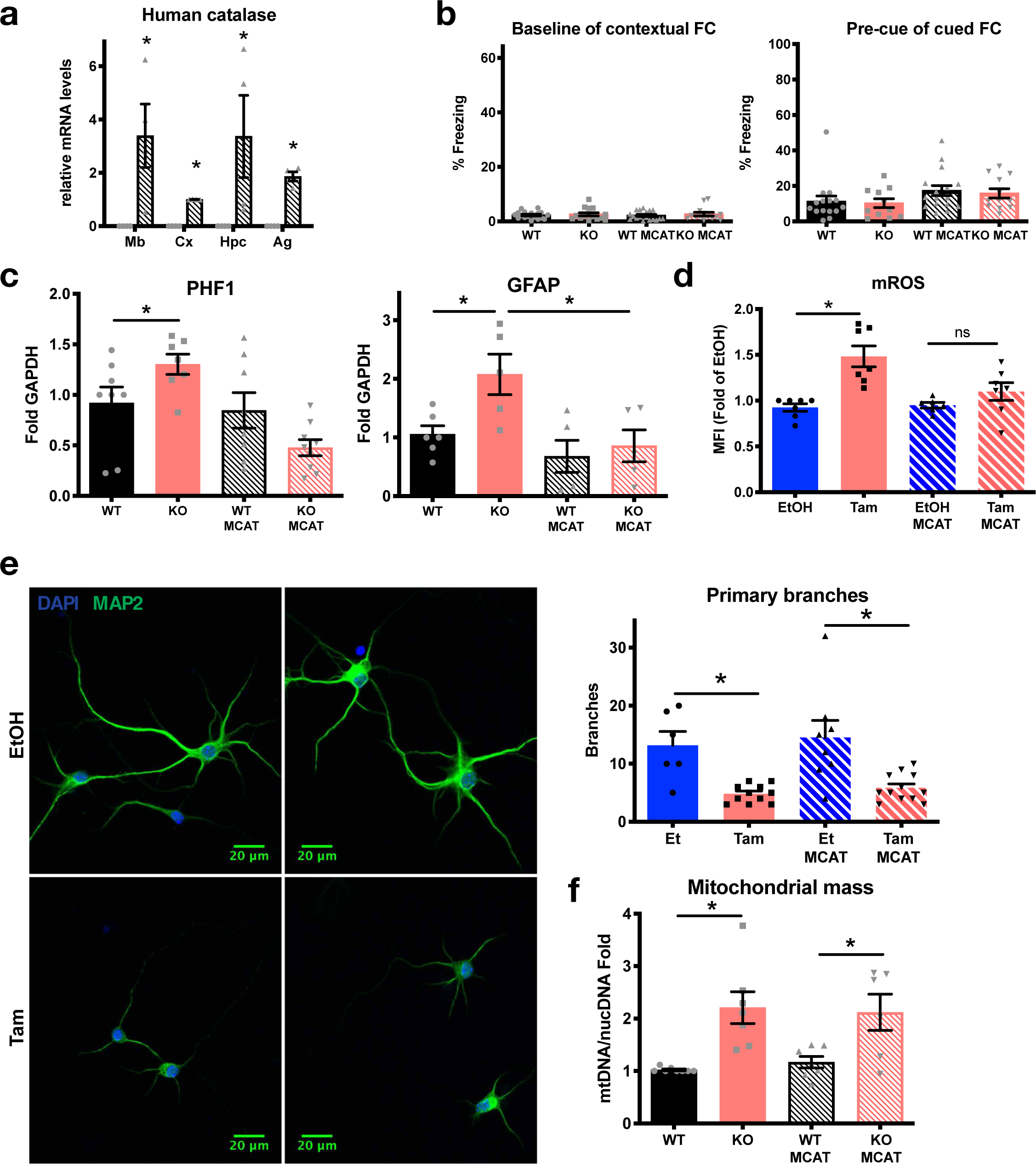
Effects of ECSIT deletion in the CNS are rescued by scavenging mROS. **a,** qPCR analysis of human MCAT expression in tissues of MCAT transgenic mouse strain (n=4 per group). *, p<0.05 in Mann-Whitney test in each brain region. **b,** Baseline after training for contextual FC and cued FC, for FC tests represented in Fig 4d. **c,** Quantification of band intensities in Western blot assessing levels of GFAP and tau PHF1 in cortex. Averages of n=8 (PHF1) and 5 (GFAP) per group are shown. *, p<0.05 in one-way ANOVA with Sidak’s post hoc test. **d,** Staining with the mitochondrial ROS probe MitoSOX, analyzed by flow cytometry in primary neurons from WT or MCAT transgenic pups with or without deletion of ECSIT with Tamoxifen *in vitro* at DIV7, as in Fig. 1b. Fold over EtOH of MFI was averaged from 5 independent experiments. *, p<0.05 in Kruskal Wallis with Dunn’s post hoc test, ns: not significant. **e,** (Left) representative image of MAP2 staining and confocal acquisition in DIV7 primary neurons. (Right) Sholl analysis with Fiji software; representative of 3 independent experiments. **f,** Mitochondrial mass assessed as in Fig. 2c, in the cortex. Data are expressed as fold increase relative to WT. Averages of 6 mice per group are shown. *, p<0.05 in Kruskal Wallis with Dunn’s post hoc test. All experiments were performed with 3- to 5-month-old mice.

**Extended Data Fig. 5.**
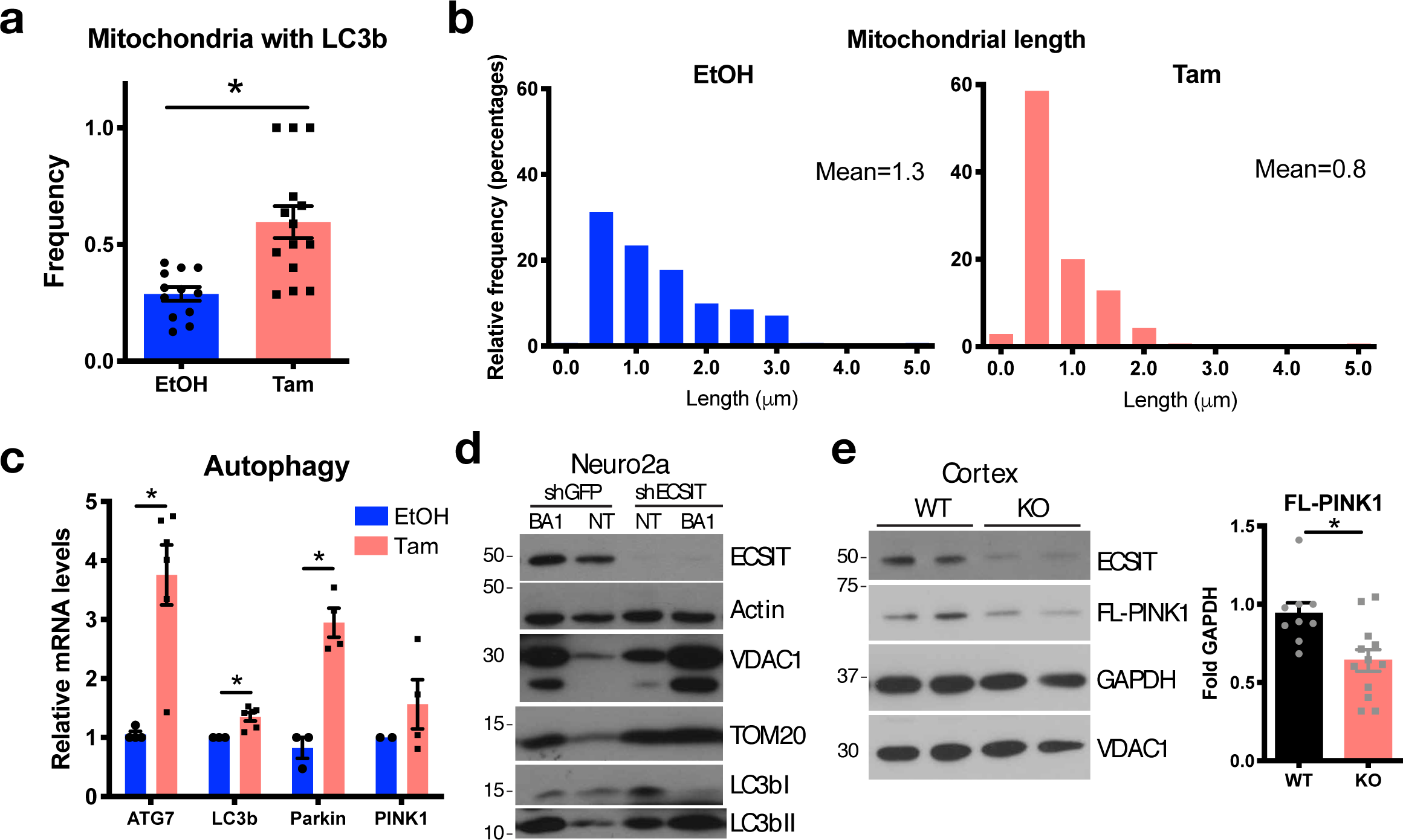
Loss of ECSIT in neurons leads to mitophagy alteration. **a,** Portion of mitochondria with enriched LC3b in primary neurons averaged from 3 independent experiments. Quantification of Fig. 5a. *, p<0.05 in Mann-Whitney test. **b,** Length of mitochondria in DIV7 primary neurons quantified in 3 independent experiments. **c,** mRNA levels of mitophagy genes in DIV7 primary neurons by qPCR. Average of 3 independent experiments, *, p<0.05 in multiple t-test. **d,** Western blot of Neuro2a mt-Keima control (shGFP) or shRNA ECSIT (shECSIT) cell lysates not treated (NT) or treated with Bafilomycin A1 (BA1) for 16 h. Representative of 4 independent experiments. **e,** Western blot analysis of full-length (FL-)PINK1 levels in cortex lysates from 3- to 5-month-old animals and quantification over GAPDH from 9 WT and 13 KO animals averaged. *, p<0.05 in Mann-Whitney test.

**Extended Data Fig. 6.**
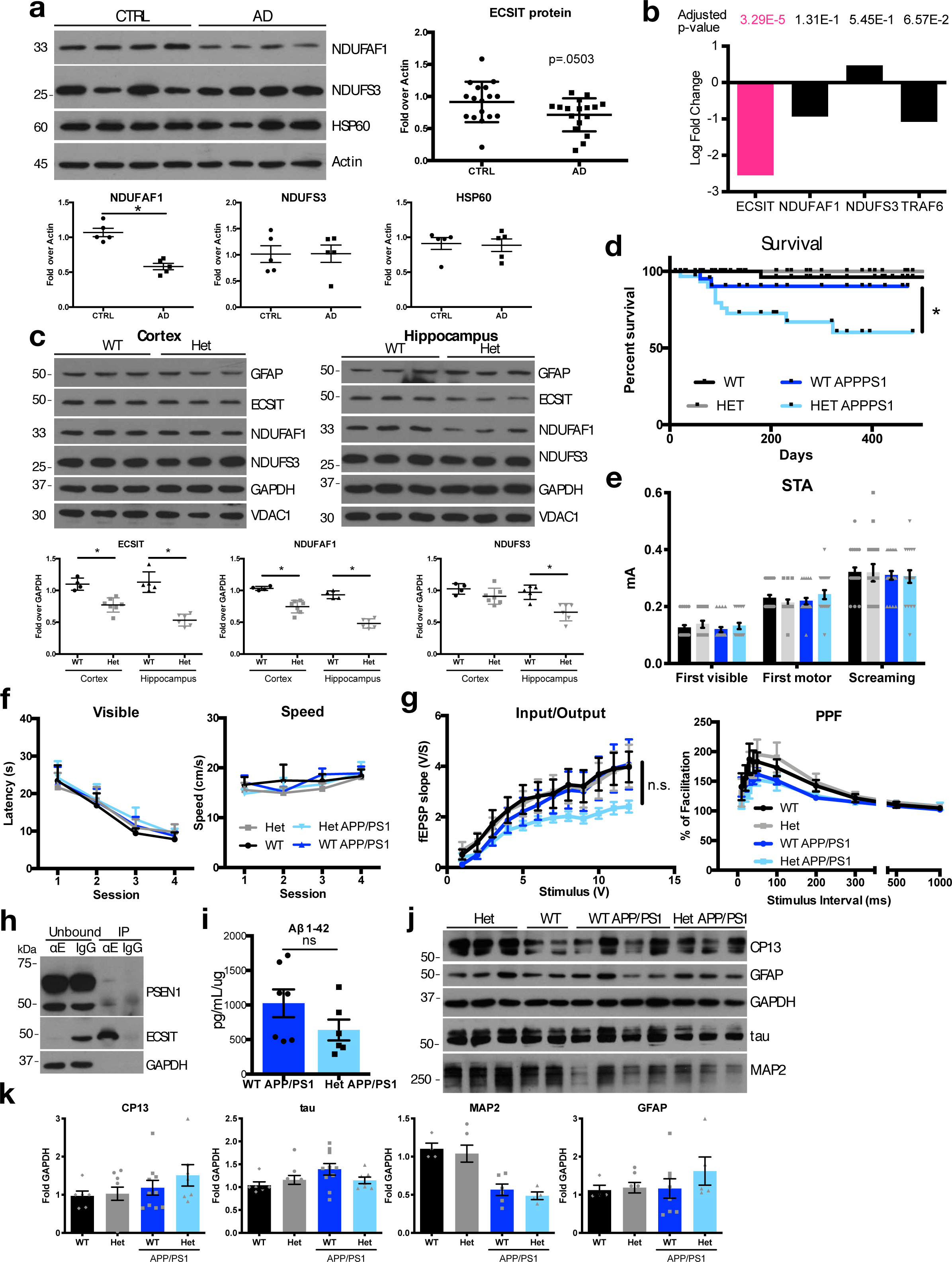
Downregulation of ECSIT expression in the context of AD contributes to disease. **a,** Western blot of complex I proteins in human control and AD patient samples and quantification of band intensities over Actin. *, p<0.05 in Mann-Whitney test. For ECSIT, data from ADRC and BBDP combined and p-value in t-test indicated. **b,** *ECSIT* mRNA levels in unaffected neurons microdissected from the entorhinal cortex of AD Braak V-VI patient samples (GSE5281). **c,** Western blot analysis of ECSIT levels in lysates of indicated brain regions Synapsin-Cre*\**E*csit* F/+ animals. Representative blots and quantification over GAPDH of 5 WT and 6 Het mice. *, p<0.05 in Mann-Whitney test in each brain region. **d,** Survival of WT (n=126) or ECSIT Het (n=73), APP/PS1 (n=21) and Het APP/PS1 (n=20) mice. *, p<0.05 in Mantel-Cox test. **e,** Sensory threshold assessment (STA) of animals tested in Fig. 6c. N=18 WT; 14 Het; 17 APP/PS1; 15 Het APP/PS1. **f,** Visual, motor and motivational controls for animals tested in Fig. 6d. N=15 WT; 10 Het; 14 APP/PS1; 12 Het APP/PS1 animals. **g,** Basal synaptic transmission and PPF in hippocampus from the animals tested in Fig. 6e (N=13 WT; 9 Het; 11 APP/PS1; 11 Het APP/PS1). ns: not significant in 2-way ANOVA and t-test for each stimulus. **h,** Coimmunoprecipitation of PSEN1 with ECSIT assessed by Western blot in adult WT mouse cortex lysates. Unbound, cell lysate supernatant after immunoprecipitation with the indicated antibodies; IP, fraction immunoprecipitated with anti-ECSIT antibody (〈 E) or IgG control. Representative of 3 independent experiments; also represented in Fig. 5f. **i**, Soluble Aβ_1-42_ ELISA in cortex of 6-month-old animals (7 WT APP/PS1 and 6 ECSIT Het APP/PS1). Not significant (ns) in t-test. **j,** Western blot analysis of indicated proteins and epitopes; quantified in **k**. **k,** Quantification of band intensities of indicated protein and epitope levels assessed by Western blotting in cortical lysates from six animals per group. Not significant in Kruskal-Wallis test. Unless otherwise specified, all experiments were performed with 3- to 5-month-old mice.

**Extended Data Fig. 7.**
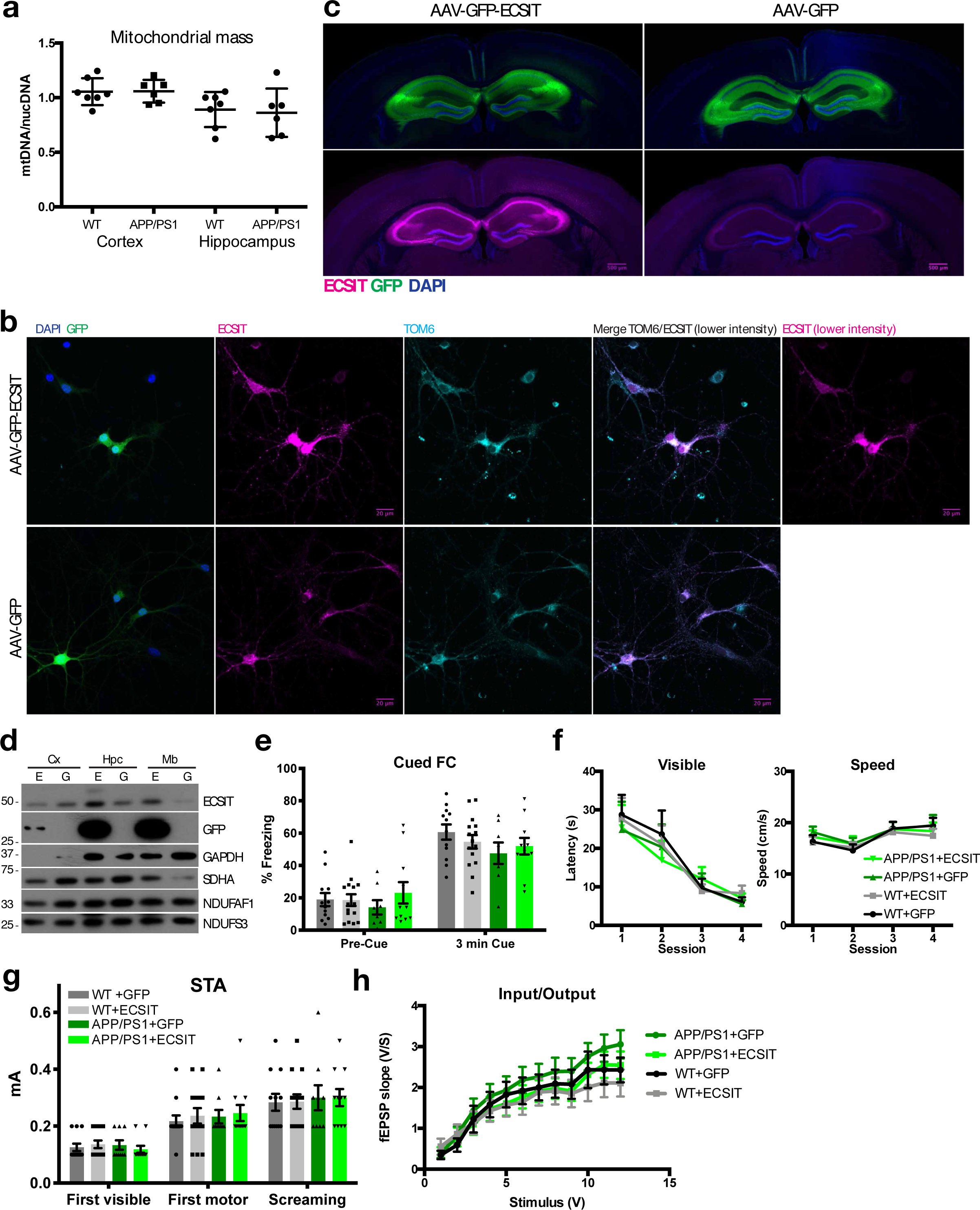
ECSIT protects against cognitive decline in AD-like pathogenesis. **a,** Mitochondrial mass assessed by qPCR. Data are expressed as fold-increase relative to WT. Average of 6 mice per group is shown. Not significant in Mann-Whitney test for each region. **b,** Representative immunofluorescence of GFP and stainings for ECSIT and TOM6 and confocal acquisition after infection of primary neurons with AAV-GFP and AAV-ECSIT-GFP. Lower intensity for ECSIT staining is shown on the left to distinguish better mitochondrial localization. Images processed with Fiji. **c,** ECSIT immunostaining and confocal imaging of coronal brain sections of mice injected with AAV-GFP-ECSIT and AAV-GFP 8 weeks after injection. Objective 10x, scale bar 500 µm. Images representative of 3 independent experiments. **d,** Western blot analysis of indicated proteins in lysates of cortex (Cx), hippocampus (Hpc) and Midbrain (Mb) of WT animals injected with AAV-GFP (G) or AAV-GFP-ECSIT (E). Representative blots of 2 experiments. **e,** Cued fear conditioning (FC) on 3- to 5-month-old WT or APP/PS1+virus animals tested in Fig. 6g. N=12 WT+GFP; 15 WT +ECSIT; 9 APP/PS1+GFP; 11 APP/PS1+ECSIT. **f,** Visual, motor and motivational controls for animals tested in Fig. 6g and h. **g,** Sensory threshold assessment (STA) of animals tested in Fig. 6g and 6h. **h,** Basal synaptic transmission in hippocampus for samples represented in Fig. 6i.

**Table S1.**
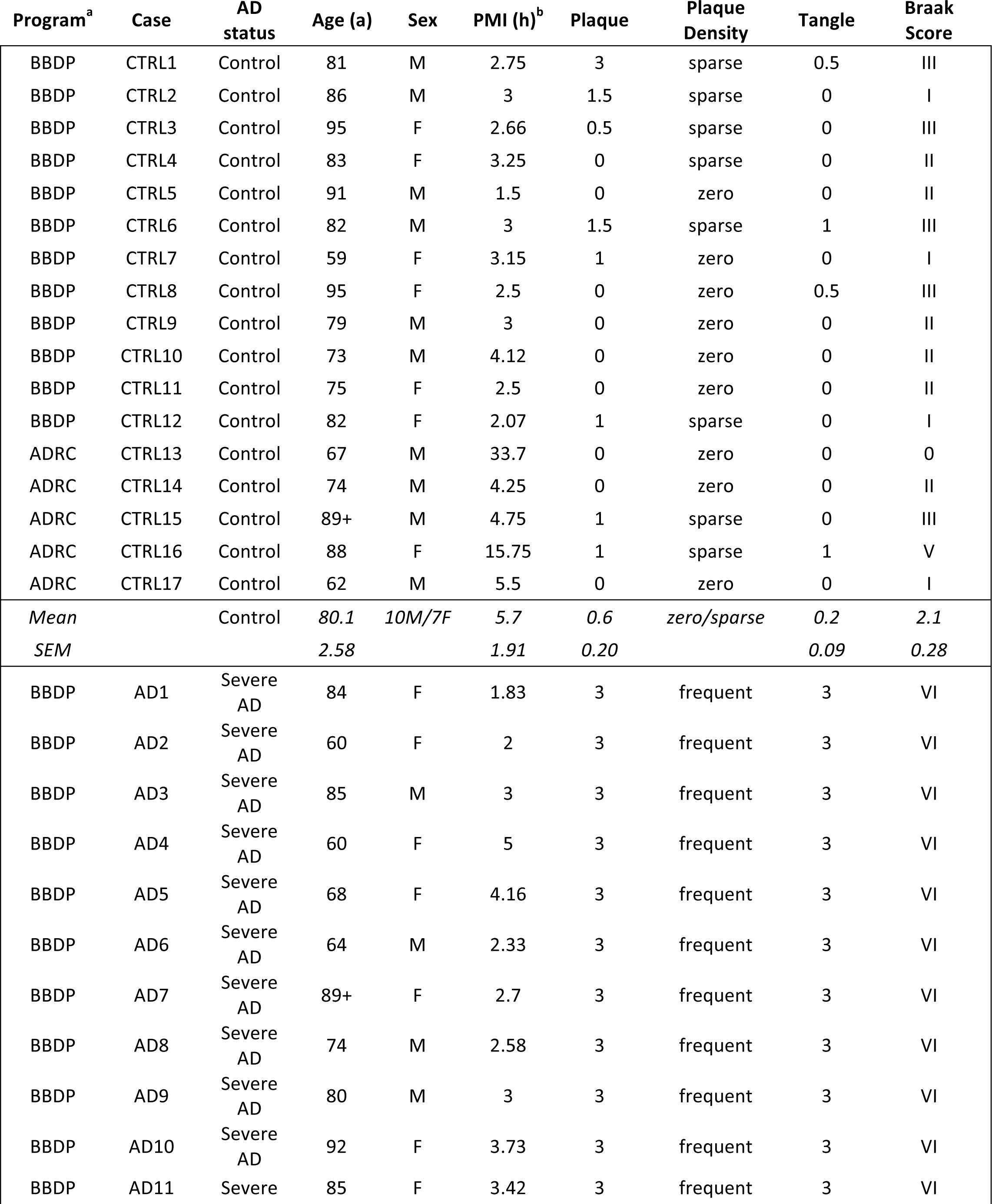
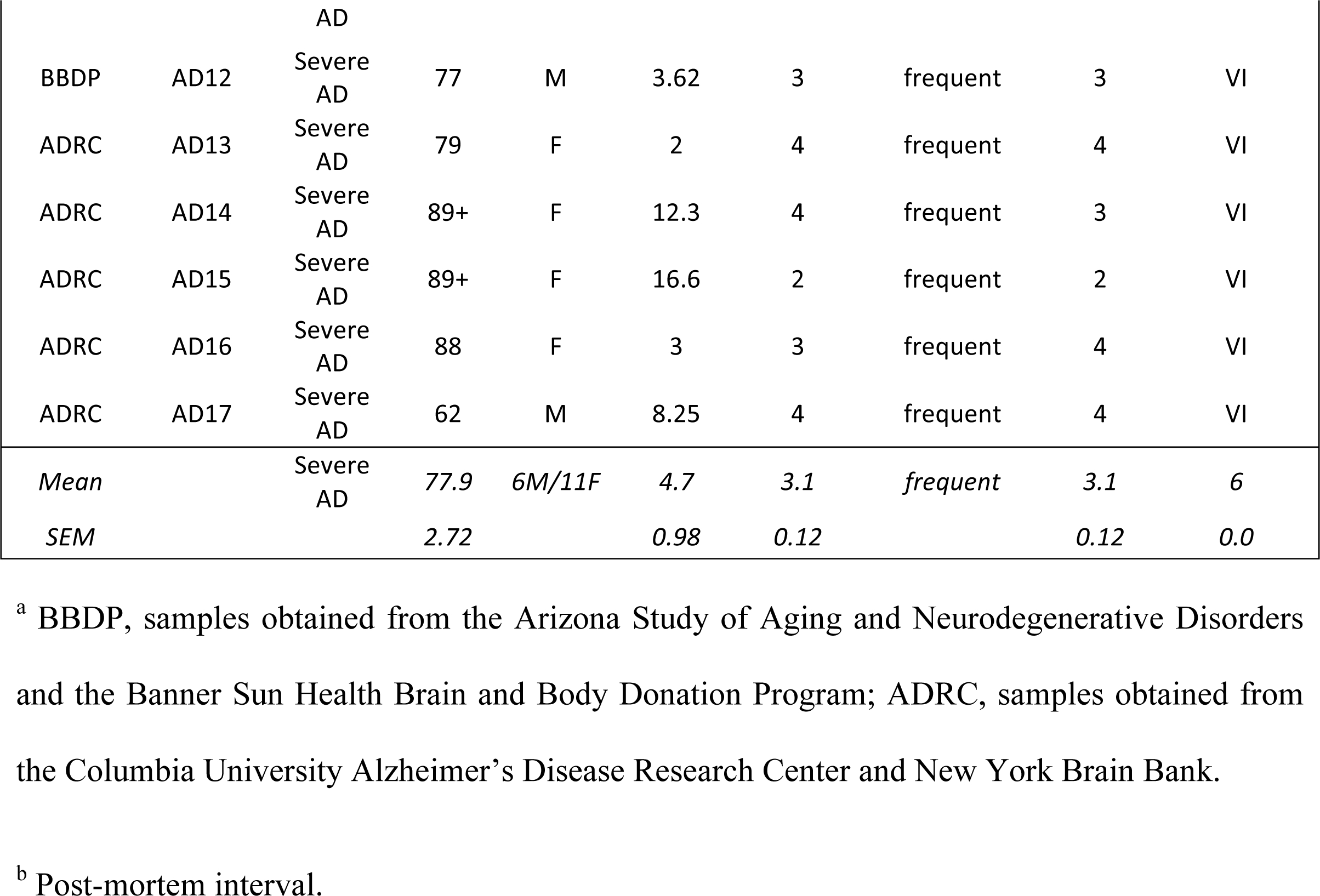
Details of human autopsy cortical tissues (temporal lobe) from individuals with Alzheimer’s disease (AD) and age-matched, non-AD controls (CTRL).

